# Chloroplast protein import determines plant proteostasis and retrograde signaling

**DOI:** 10.1101/2022.03.19.484971

**Authors:** Ernesto Llamas, Seda Koyuncu, Hyun Ju Lee, Ricardo Gutierrez-Garcia, Nick Dunken, Nyasha Charura, Salvador Torres-Montilla, Elena Schlimgen, Pablo Pulido, Manuel Rodriguez-Concepcion, Alga Zuccaro, David Vilchez

## Abstract

Proteins containing polyglutamine (polyQ) repeats are prone to aggregation and can lead to distinct human pathologies. For instance, Huntington’s disease is caused by an abnormal expansion of the polyQ stretch (> Q35) of Huntingtin (HTT) protein. However, plants express hundreds of proteins containing polyQ regions, but no pathologies arising from these factors have been reported to date. Here, we ask how plants maintain the proteostasis of polyQ-containing proteins, which are intrinsically enriched in the plant proteomes. To this end, we overexpressed an aggregation-prone fragment of human HTT (Q69) in plant cells. In contrast to invertebrate and mammalian transgenic models, we find that *Arabidopsis thaliana* plants suppress Q69 aggregation. This elevated proteostasis ability is mediated through the import and degradation of Q69 in chloroplasts. Conversely, inhibition of chloroplast protein import either genetically or pharmacologically reduces the capacity of plant cells to prevent Q69 aggregation. We find that Q69 interacts with the chloroplast stromal processing peptidase (SPP). Notably, expression of synthetic Arabidopsis SPP is sufficient to suppress aggregation of polyQ-expanded HTT in human cells. Beyond ectopically expressed Q69-HTT, endogenous polyQ-containing proteins also aggregate in Arabidopsis upon inhibition of chloroplast import. Among them, the plastid casein kinase 2 (pCK2), which contains a polyQ region next to the chloroplast targeting sequence motif, can also be localized into the nucleus. Upon inhibition of chloroplast import, pCK2 accumulates at higher levels in the nucleus and forms diamond-shaped amyloid-like fibrils surrounding the chloroplasts. These results indicate that the differential conformation and redistribution of pCK2 to the nucleus depends on chloroplast import efficiency, providing a role of polyQ repeats in chloroplast to nucleus communication (*i*.*e*. retrograde signaling). Together, our findings establish chloroplast protein import and proteases as determinants of polyQ proteostasis, with important implications for plant biology that can also lead to therapeutic approaches for human diseases that involve protein aggregation.

## Introduction

Native proteins must be correctly folded, assembled with their binding partners, and targeted to their appropriate subcellular destinations (Braselmann et al. 2013). However, conformational changes in specific proteins can result in their spontaneous self-assembly, forming toxic aggregates that cause distinct human neurodegenerative diseases. For example, the prion protein (PrP) can misfold and aggregate, leading to transmissible neurodegenerative disorders (Martinelli et al. 2019). Most neurodegenerative diseases are associated with proteins containing prion-forming domains (PFDs) or intrinsically disordered regions (IDRs) rich in asparagine (N) and glutamine (Q) residues that can drive aggregation (Ross et al. 2004). For instance, a common feature of polyQ-containing proteins is the capacity to convert into a priogenic form, and these self-replicating conformations mediate pathogenic and nonpathogenic processes in yeast and higher eukaryotes (Shorter and Lindquist 2005). Cells employ distinct proteostasis pathways to avoid the hazardous accumulation of polyQ expanded proteins, including degradation through the ubiquitin-proteasome system (UPS) and disaggregation through members of the Heat shock protein (Hsp) family (Nath and Lieberman 2017; Kitamura et al. 2006; Gruber et al. 2018; Doi et al. 2004; Braun et al. 2010; Koyuncu et al. 2018; Noormohammadi et al. 2016). At least nine human neurodegenerative diseases are associated with proteins containing polyQ repeats. Among them, Huntington’s disease (HD) is caused by mutations in the *huntingtin* (*HTT*) gene (Finkbeiner 2011; Koyuncu et al. 2017). The wild-type HTT protein contains 6–35 polyQ repeats (Saudou and Humbert 2016; Cattaneo et al. 2005), and does not aggregate even under stress conditions (Koyuncu et al. 2018). In individuals affected with HD, mutations expand the CAG triplet repeat region in the exon 1, resulting in an unstable expanded polyQ stretch (> Q35) that causes aggregation and proteotoxicity. Expression of the pathogenic fragment of the exon 1 of the mutant polyQ-expanded HTT in different model organisms is sufficient to recapitulate key aspects of HD, including pathological protein aggregation and cell death (Pearce and Kopito 2018). Another human disease-related protein, ATXN3, can contain up to 52 polyQ repeats in its wild-type form, but still does not form aggregates even under challenging conditions (Koyuncu et al. 2018). However, a polyQ extension beyond 52 repeats triggers ATXN3 aggregation, causing Machado-Joseph disease (Ranum et al. 1995; Kawaguchi et al. 1994).

In contrast to human proteins such as HTT and ATXN3 that have relatively long polyQ stretches in their wild-type forms, the polyQ repeats never exceed 24 polyQ repeats in the *Arabidopsis thaliana* proteome (Kottenhagen et al. 2012). Interestingly, a few prion-like proteins with polyQ tracts were recently characterized to function as sensors that integrate internal and external cues, allowing Arabidopsis to adapt to ever-changing environmental conditions (Jung et al. 2020; Dorone et al. 2021; Chakrabortee et al. 2016). For instance, the transcription factor EARLY FLOWERING 3 (ELF3) contains a Q7 stretch, which enables the plant to respond to high temperatures. As such, the Q7 stretch is necessary and sufficient to make ELF3 adopt two reversibly conformations. At 22 ºC, ELF3 is soluble and binds genes that repress flowering. At temperatures higher than 27 ºC, ELF3 forms aggregates that relieve transcriptional repression and promote flowering (Jung et al. 2020; Alberti 2020). Thus, Arabidopsis ELF3 with a relative short Q7 motif can form aggregates under stress conditions (Jung et al. 2020).

To shed light on plant proteostasis mechanisms regulating polyQ prion-like proteins, we expressed the exon 1 of the human HTT gene encoding an aggregation-prone polyQ stretch. Since the maximum polyQ expansion in plant proteins is 24 repeats (Kottenhagen et al. 2012), we generated Q28 and Q69 models to examine how plants can cope with abnormal polyQ proteins. Notably, stably transformed Arabidopsis plants expressing either Q28 or Q69 in the cytosol did not exhibit aggregates or deleterious phenotypes under normal conditions. However, similar to the Arabidopsis ELF3 (Q7) (Jung et al. 2020), we observed a heat-dependent aggregation of polyQ proteins in stable transgenic plants expressing either Q28 or Q69. Under normal conditions, Arabidopsis plants prevented protein misfolding and aggregation of polyQ-expanded proteins through their import and degradation into the chloroplast. Protein aggregation and cleavage assays using human cells as a heterologous system indicated that the stromal processing peptidase (SPP) is a key chloroplast factor to reduce aggregation of polyQ-extended proteins. Moreover, we unveiled that genetical and pharmacological impairment of chloroplast import triggers the cytosolic accumulation and aggregation of Q69. In addition to ectopic Q69, inhibition of chloroplast import also results in the aggregation of endogenous Arabidopsis prion-like proteins containing polyQ stretches. Among them, we focused on the plastid casein kinase 2 (pCK2), a factor involved in chloroplast-to-nucleus communication (Wang et al. 2014). pCK2 was reported as a chloroplast localized protein (Salinas et al. 2006; Mulekar and Huq 2015) and we found that it contains a polyQ prion-like domain next to its chloroplast targeting sequence motif. Surprisingly, pCK2 accumulated in the nucleus and formed diamond-shaped fibril amyloids surrounding chloroplasts upon inhibition of chloroplast protein import. The pCK2 redistribution to the nucleus upon impaired chloroplast import reveals its uncharacterized role in retrograde signaling. Moreover, our work opens a new door for the discovery of therapeutic targets against human polyQ diseases.

## Results

### Constitutive and stable expression of Q69 in Arabidopsis does not cause protein aggregation under normal conditions

In multiple invertebrate and mammalian model organisms, overexpressing the HTT exon 1 containing more than 35 glutamine repeats is sufficient to trigger protein aggregation (Mangiarini et al. 1996; Morley et al. 2002; Gruber et al. 2018). To recapitulate the pathological aggregation phenotype of HD in plants, we generated transgenic plants overexpressing the human mutant HTT exon 1 fragment. To this end, we generated the constructs *35S:Citrine-HTTexon1-Q28* (Q28) and *35S:Citrine-HTTexon1-Q69* (Q69) **(Fig. 1a)**. We first analyzed the distribution of Q28 and Q69 proteins in *Nicotiana benthamiana* leaf cells by agroinfiltration of the corresponding constructs. As expected, transient Q69 (tQ69) overexpression resulted in the formation of densely packed amorphous aggregates **(Fig. 1b)**. Interestingly, tQ69 aggregates were found in some cells while others showed a homogenous distribution, suggesting that proteostasis collapse varies between cells **(Fig. 1b and Supplementary Fig. 1)**. In contrast, transiently expressed Q28 (tQ28) was diffusely distributed in the cytoplasm and the nucleus even after several days post-agroinfiltration **(Fig. 1b)**. At 12 days post-agroinfiltration, *N. benthamiana* leaves expressing tQ69 exhibited chlorosis, an indicator of cell death. However, we did not observe this phenotype in agroinfiltrated leaves expressing tQ28 **(Fig. 1c)**. Together, our data indicate that transient acute overexpression of HTTexon1-Q69 collapses the proteostasis network in mature *N. benthamiana* leaves.

**Figure 1.**
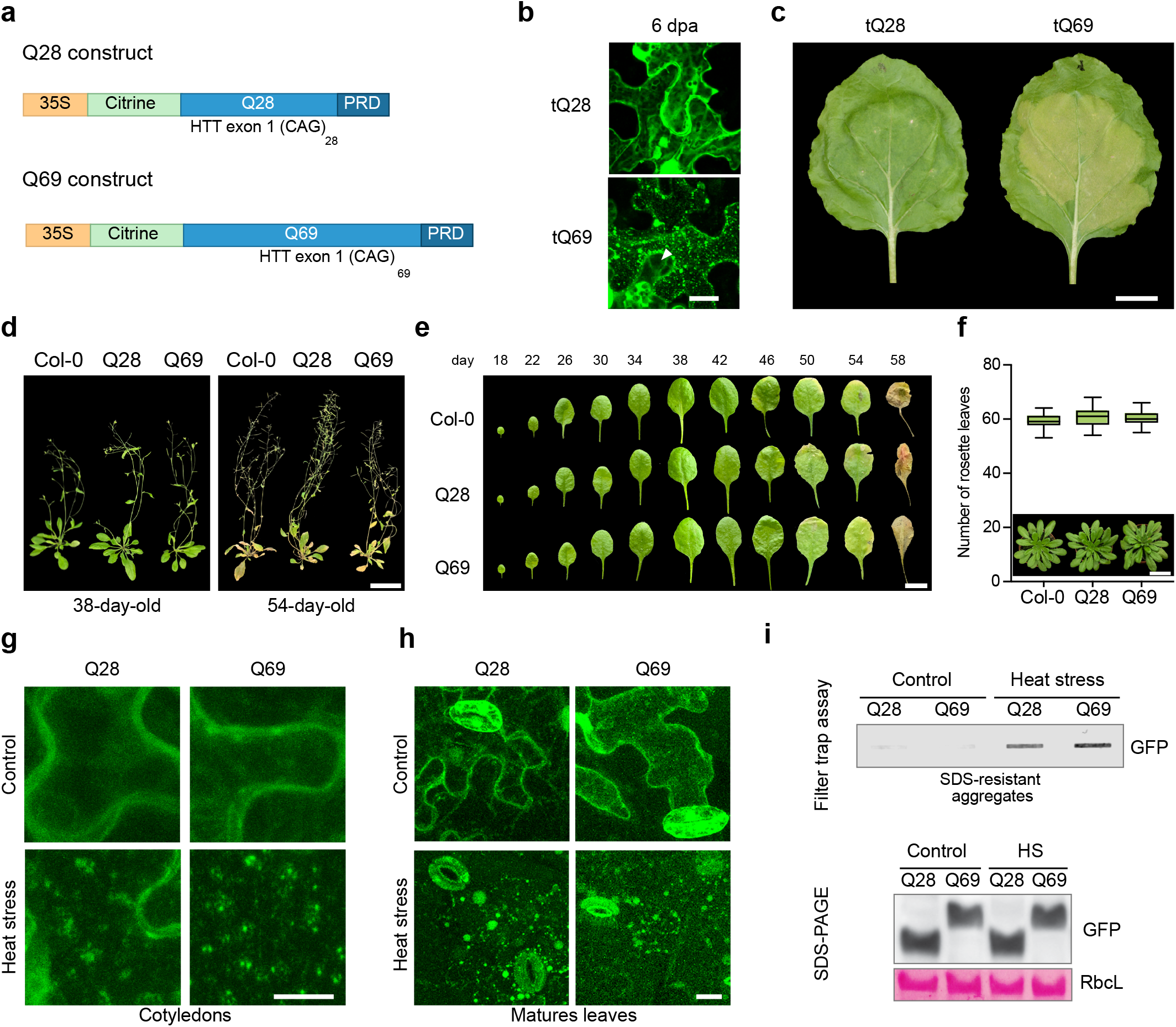
Plants constitutively expressing polyQ proteins do not display aggregates or deleterious effects under normal growth conditions. **(a)** Schematic representation of the constitutive constructs Q28 and Q69. Proline-rich domain (PRD). **(b)** Q28 and 69 distribution in *N. benthamiana* epidermal pavement cells from leaves analyzed at 6 days post-agroinfiltration (dpa). Citrine fluorescence (Citrine is a fluorescent protein derived from GFP) and bright field are shown. Scale indicates 20 μm. **(c)** Phenotype of 6-week-old *N. benthamiana* leaves after 12 dpa with Q28 or Q69. Scale indicates 2 cm. **(d)** Phenotype of mature and senesce Col-0, Q28, and Q69 plants. Scale represents 5 cm. **(e)** Life span leaf analysis of the fourth true leaf from stable Arabidopsis Q28 and Q69 compared to Col-0. Scale indicates 1 cm. **(f)** Flowering time of polyQ and Col-0 plants grown under short-day conditions at 22 ºC. Representative images of 46-day old plants are shown. Scale indicates 5 cm. The statistical comparisons between Col-0 and polyQ plants were made by two-tailed Student’s t-test for unpaired samples where not significant (P > 0.05) differences were found. **(g)** Confocal images showing Q28 and Q69 distribution in epidermal pavement cells from cotyledons. 7-day-old seedlings grown at 22 ºC were transfer in dark to incubators at 45 ºC (heat stress) or 22 ºC (control) for 90 minutes. Scale indicates 10 μm. **(h)** Q28 and Q69 distribution in epidermal pavement cells from leaves of 22-day-old plants. The 4th leave of Q28 or Q69 plants were dissected and incubated in dark under heat stress or control conditions for 90 minutes. Scale indicates 25 μm. **(i)** Filter trap and SDS-PAGE analysis with antibody against GFP (capable of recognizing Citrine tag) of Q28 and Q69 seedlings used for microscopy analysis in d. RbcL is shown as loading control. The images are representative of two independent experiments.

Next, we generated and characterized stable Arabidopsis transgenic plants expressing Q28 and Q69 under the control of the 35S promoter **(Supplementary Fig. 2)**. To our surprise, constitutive overexpression of Q69 did not cause deleterious effects in Arabidopsis plants **(Fig. 1d-f)**. As such, transgenic Q28 and Q69 lines had similar development, life span, flowering time, and photosynthetic activity to untransformed Col-0 wild-type (WT) controls **(Fig. 1d-f and Supplementary Fig. 2)**. Notably, we observed that both Q28 and Q69 proteins were distributed in a diffuse pattern among the root tips, cotyledons, and mature leaves of these transgenic plants when growing under normal conditions **(Supplementary Fig. 2)**. Additionally, proteostasis stress markers indicated absence of folding stress in these polyQ lines **(Supplementary Fig. 2)**. Taken together, our results indicate that when stable expression of expanded Q69 proteins occurs from early stages of development, Arabidopsis plants have mechanisms to sustain proteostasis and prevent the accumulation of polyQ aggregates throughout plant life.

In humans, wild-type HTT protein ranges between 6-35 polyQ repeats and other proteins such as ATXN3 can contain up to 52 Q repeats before becoming prone to aggregation (Ranum et al. 1995; Kawaguchi et al. 1994). In contrast, the polyQ stretches in endogenous Arabidopsis proteins do not exceed 24 Q repeats (Kottenhagen et al. 2012) **(Supplementary Data 1)**. Among them, a recent study demonstrated that the Arabidopsis transcription factor ELF3 localizes diffusely in the cell at 22 ºC. However, ELF3 forms aggregates at higher temperatures even when it only contains a short polyQ7 stretch (Jung et al. 2020). Thus, we hypothesized that in contrast to animal models (Morley et al. 2002; Brignull et al. 2006), relatively shorter polyQ stretches are prone to aggregation in plants during stress conditions. Thus, plants may employ intrinsic proteostasis mechanisms to avoid polyQ aggregation under normal conditions. To study the possible alternation of conformational states of polyQ-extended proteins at elevated temperatures, we treated 7-day-old stable transgenic plants Q28 and Q69 with mild (37 ºC) and severe heat stress (45 ºC) for 90 minutes. Although mild stress conditions did not cause aggregation of cytosolic Q28 and Q69 **(Supplementary Fig. 3)**, a severe heat stress led to the formation of Q28 and Q69 aggregates **(Fig. 1g-i)**. However, Q28 and Q69 seedlings did not show increased sensitivity to heat stress treatment compared to wild-type plants **(Supplementary Fig. 4)**.

### Q28 and Q69 proteins interact with several chloroplast proteostasis components

To determine the mechanisms underlying the enhanced ability of plants to prevent polyQ aggregation under control conditions, we examined the interactome of Q28 and Q69 in Arabidopsis plants. To this end, we performed co-immunoprecipitation (co-IP) experiments followed by a single shot Cotyledons Matures leaves label-free proteomic approach. Pulldown experiments indicated that Q28 and Q69 were the most enriched proteins **(Fig. 2 and Supplementary Data 2)**. Hierarchical clustering revealed a similar protein interaction between Q69 and Q28 lines **(Supplementary Fig. 5)**. The proteomics data suggest that plant proteostasis interactors do not differ between Q28 and Q69, since Q24 is the longest polyQ stretch in Arabidopsis endogenous proteins.

**Figure 2.**
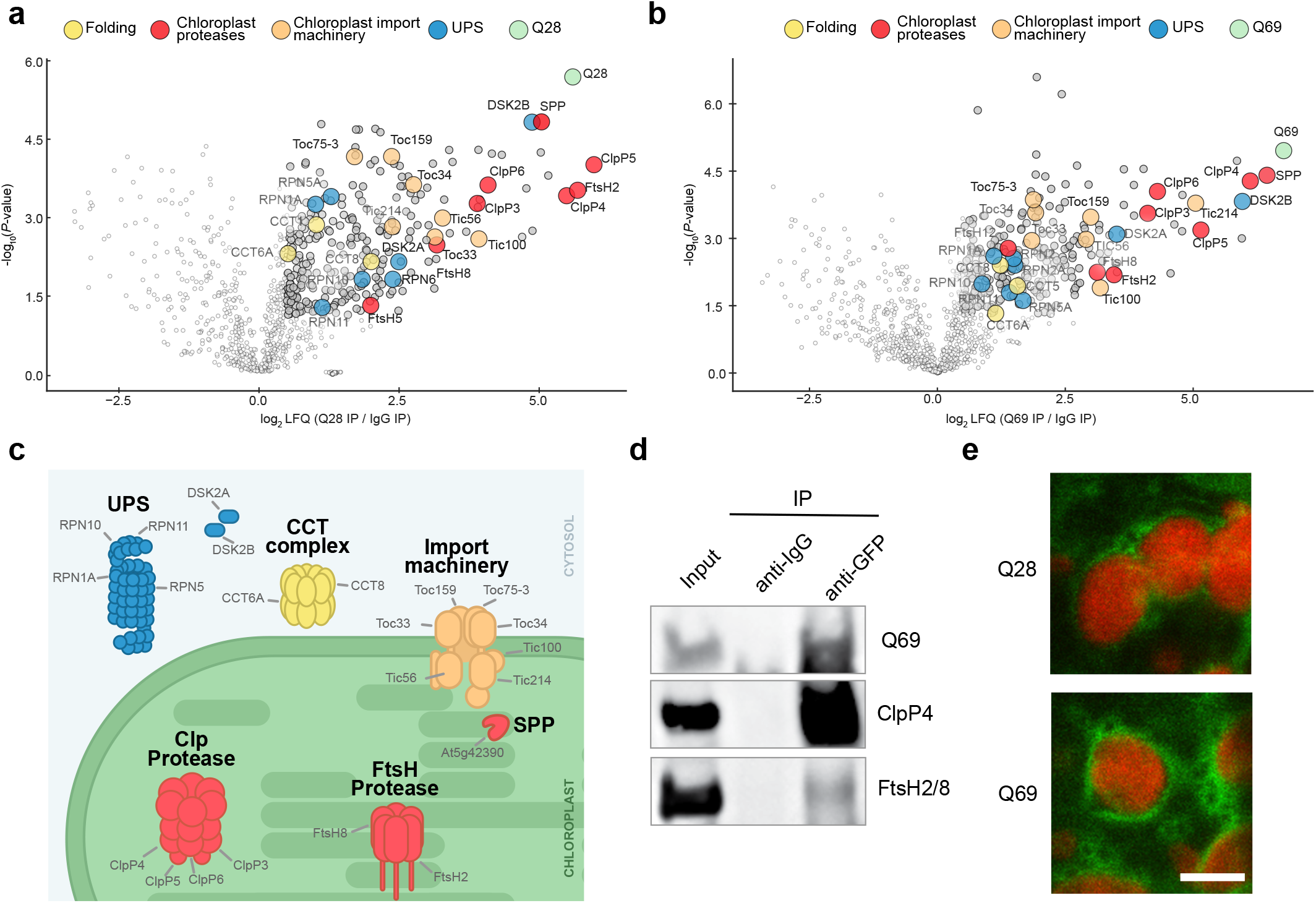
Q28 and Q69 proteins interact with cytosolic and chloroplast proteostasis components. **(a)** Volcano plot of Q28 and Q69 **(b)** interactome in transgenic Arabidopsis 7-day-old seedlings (n = 3). -log_10_(P-value) of a two-tailed t-test is plotted against the log_2_ ratio of protein label-free quantification (LFQ) values from co-IP experiments using GFP antibody (against Citrine-Q28 or Citrine-Q69) to control co-IP with IgG antibody. Gray and colored circles indicate significance after correction for multiple testing (false discovery rate (FDR) < 0.05 was considered significant). Yellow colored circles represent proteins involved in protein folding, red colored proteins involved in chloroplast proteolytic degradation, in orange proteins that form part of the chloroplast import machinery, in blue proteins involved in the UPS, and green circles represent Q28 or Q69 proteins. **(c)** Scheme indicating the subcellular localization of selected common interactors of Q69 and Q28. **(d)** Co-IP with GFP and control IgG antibodies in Q69 seedlings followed by immunoblot against the chloroplast protease subunits ClpP4 and FtsH2/8. Images are representative of three independent experiments. **(e)** Q28 and Q69 distribution in mesophyll cells of 7-day-old cotyledons. Images show Citrine fluorescence (green) and chloroplast autofluorescence (red). Scale indicates 5 μm.

Among the proteins interacting with Q28 and Q69, we found several involved in cytosolic protein folding and the UPS **(Fig. 2a-c and Supplementary Data 2)**. Interestingly, we detected the ubiquitin binding receptor proteins DSK2A and DSK2B as Q28 and Q69 interactors in Arabidopsis **(Fig. 2a-c)**. Importantly, DSK2 (also known as Ubiquilin) suppresses polyQ-induced protein aggregation and toxicity in animal models of Huntington’s disease (Wang et al. 2006). Moreover, we also identified subunits of the chaperonin TRiC/CCT as polyQ interactors **(Fig. 2a-c)**. The TRiC/CCT complex reduces the accumulation of polyQ aggregates in human cells (Noormohammadi et al. 2016). Similar to other interactome studies of the mutant HTT exon 1 expressed in mammalian cells (Kim et al. 2016), we identified several proteasome subunits as interactors of polyQ protein in plants **(Fig. 2a-c)**. In animal cells, polyQ-extended proteins accumulate as aggregates when proteasome activity is impaired (Koyuncu et al. 2018). Similarly, transferring Q69 plants to media supplemented with the proteasome inhibitor MG-132 resulted in the formation of aggregates **(Supplementary Fig. 6)**.

In addition to cytosolic components of the proteostasis network, our interactome analysis revealed that polyQ proteins also bind to chloroplast proteins, including import machinery components and proteases such as Clp and FtsH **(Fig 2a-d)**. Moreover, confocal microscopy analyses indicated that Q28 and Q69 localize around the chloroplasts **(Fig. 2e and Supplementary Fig. 7)**. Together, our proteomics analyses suggest a link between chloroplasts and proteostasis of polyQ-containing proteins.

### Disruption of chloroplast proteostasis causes accumulation and aggregation of Q69 in the cytosol

Notably, the interactor partners of both Q28 and Q69 are enriched for factors involved in protein importing and protease-mediated degradation pathways in the chloroplast **(Fig. 2a-c)**. Most chloroplast proteins are encoded by the nuclear genome, synthesized in the cytosol, and then imported into such organelle through the TIC/TOC machinery as unfolded chloroplast protein precursors (or pre-proteins). Pre-proteins contain an unstructured/unfolded N-terminal transit peptide (Lee and Hwang 2021; Lee et al. 2013), which is recognized by the TIC/ TOC complex and transported into the stroma to be subjected to proteolytic processing by several proteases (Sun et al. 2021; Thomson et al. 2020).

The unstructured nature of the Q69 protein led us to hypothesize that it can be recognized as a pre-protein by the chloroplast import machinery. To assess whether endogenous Arabidopsis proteins that are targeted to the chloroplast contain polyQ domains, we analyzed the Arabidopsis proteome searching for polyQ stretches in annotated chloroplast proteins **(Supplementary Data 1)**. We found that 5 out of 6 nucleus encoded-chloroplast proteins have the polyQ stretches close to the N-terminal chloroplast transit peptide **(Supplementary Fig. 8)**. Prediction software indicated that the polyQ-stretches from chloroplast proteins are embedded in prion-like domains (PrLDs) or intrinsically disordered regions (IDRs) **(Supplementary Fig. 8)**. Likewise, the Q69 protein, which has a large prion-like/disorder domain, was also predicted to be a chloroplast protein **(Supplementary Fig 8)**. To assess the possible localization of Q69 into the chloroplast, we used lincomycin (LIN) to impair chloroplast protein import and Clp protease-mediated degradation (Llamas et al. 2017; Wu et al. 2019). Transfer of 7-day-old Q69 seedlings to liquid media supplemented with 800 µM LIN rapidly caused the accumulation and aggregation of Q69 **(Fig. 3a, b)**. Notably, after 24 hours of treatment with LIN, Q69 remained aggregated but its total levels were reduced **(Fig. 3a-c)**. These results suggest that when chloroplast import is impaired, proteasome activity increases as a compensatory mechanism to maintain cytosolic proteostasis by degrading excess of misfolded Q69. Treatment with lower concentrations of LIN (15 µM) allowed detection of Citrine fluorescence within some chloroplasts **(Fig. 3c)**, suggesting that Q69 is imported and degraded in the chloroplast. In addition, we expressed Q69 in *toc159*, a mutant line with altered chloroplast import. Similar to LIN treatment, Q69 protein aggregates in *toc159* mutant plants **(Fig. 3e, f)**. However, we did not observe higher total Q69 levels in long-term LIN treated plants or in *toc159* mutant plants compared to their respective controls **(Fig. 3f and Supplementary Fig. 9)**. Prolonged import defects may trigger Q69 degradation through the proteasome to reduce the formation of cytosolic aggregates, a compensatory mechanism similar to that observed when chloroplast pre-proteins accumulate in the cytosol (Thomson et al. 2020; Wu et al. 2019). Collectively, our data suggest that misfolded versions of Q69 are targeted to the chloroplasts for further degradation under normal conditions, hence avoiding the accumulation of aggregates in the cytosol. On the other hand, when chloroplast import is transiently impaired, cytosolic Q69 levels rapidly increase surpassing a threshold that triggers the formation of aggregates.

**Figure 3.**
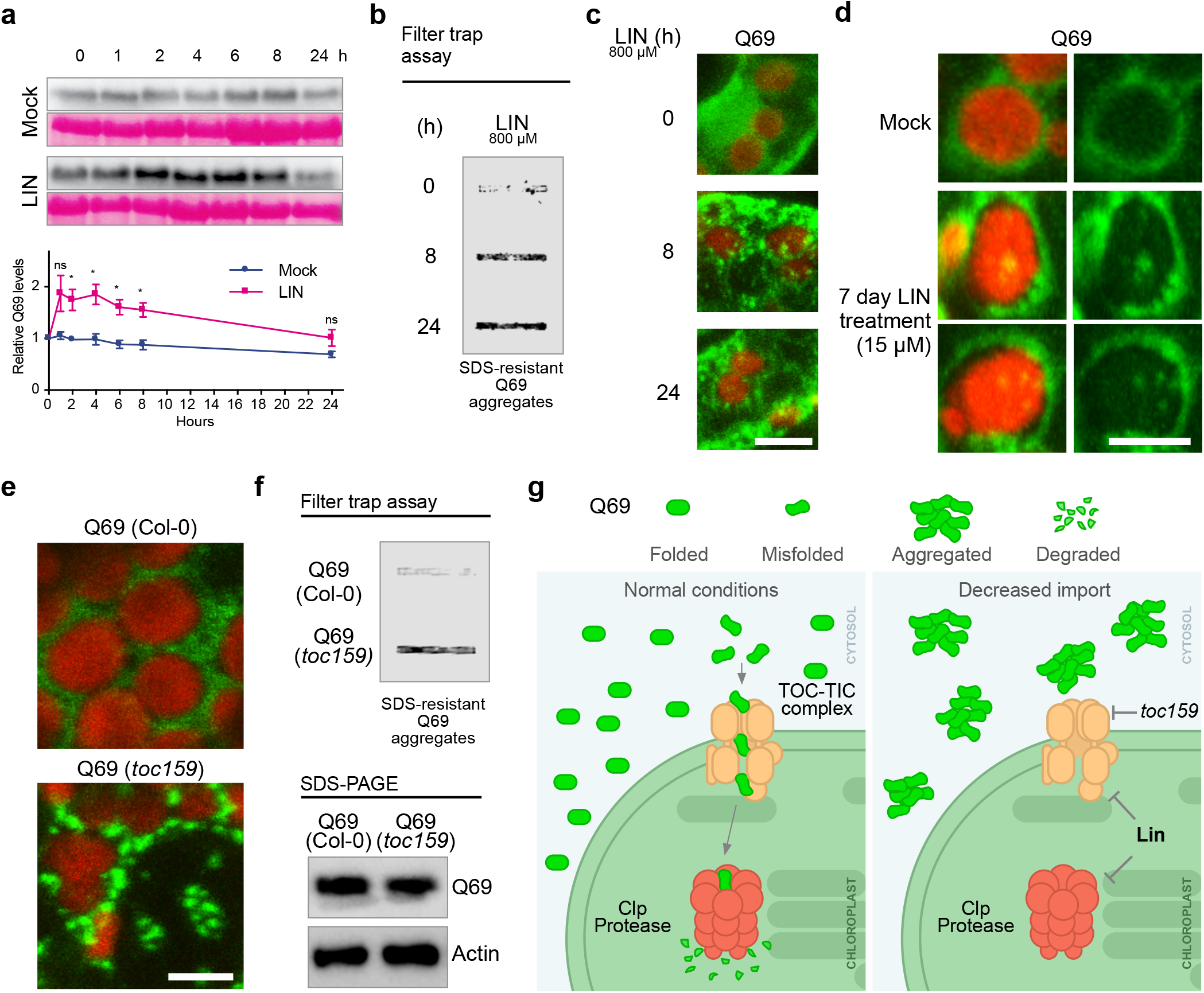
Reduced chloroplast protein import triggers aggregation of Q69. **(a)** Immunoblot analyses of 7-day-old Q69 plants transfer to liquid MS supplemented with mock or LIN. RbcL was used as loading control. Representative immunoblots of 4 independent experiments. Lower graphs show relative Q69 protein levels after treatment with mock and LIN. Chart represents the mean ± s.e.m of 4 independent experiments. The statistical comparisons between mock- and LIN-treated samples were made by two-tailed Student’s t-test for unpaired samples. P value: *P < 0.05, **P < 0.01, ns, not significant. **(b)** Filter trap analysis showing Q69 accumulation and aggregation after LIN treatment. 3 independent filter traps were performed showing similar results. **(c)** Representative images showing Q69 aggregation in stomata from cotyledons after LIN treatment. Images show Citrine (green) and chloroplast autofluorescence (red). Scale indicates 5 μm. **(d)** Representative images showing detection of Citrine fluorescence within the chloroplast of epidermal hypocotyl cells. Images show Citrine (green) and chloroplast autofluorescence (red). Scale indicates 5 μm. **(e)** Representative images showing Q69 distribution in Col-0 and *toc159* background. 7-day-old seedlings were analyzed. Images taken in mesophyll cells. Images show Citrine (green) and chloroplast autofluorescence (red). Scale indicates 5 μm. **(f)** Filter trap and SDS-PAGE analysis of samples indicated in e. Actin was used as loading control. Results from 2 independent experiments are shown. **(g)** Proposed model of how chloroplast modulates Q69 status. During normal conditions (right panel), Q69 distributes homogenously surrounding the chloroplasts, while misfolded/unstructured versions are imported to the chloroplast to further degradation. When protein import or chloroplast degradation is impaired using LIN (left panel), misfolded Q69 accumulates in the cytosol promoting the formation of Q69 aggregates.

### The chloroplast stromal processing peptidase (SPP) cleaves and reduces aggregation of polyQ expanded proteins

Given that Q69 can be imported to the chloroplast for degradation, we aimed to unveil possible plastidial degradation mechanisms. Our proteomics data revealed that both Q28 and Q69 interact with the stromal processing peptidase (SPP) **(Fig. 2a, b)**. SPP initially recognizes pre-proteins, binds to their transit peptide, removes it through a single endoproteolytic step, and then cleaves the released transit peptide to subfragmented forms (Zhong et al. 2003). We hypothesized that SPP might recognize and cleave imported unstructured Q69, hence reducing proteotoxicity in the chloroplast. To test this hypothesis, we co-transfected HEK293 cells with mRFP-HTTexon1-Q74 (mRFP-Q74) and a synthetic SPP (without chloroplast transit peptide and codon optimized) fused to GFP in the N-terminal (GFP-SPP) **(Fig. 4a, b)**. When we co-expressed mRFP-Q74 and GFP-SPP, we observed a reduction of aggregation and cleavage of mRFP-Q74 compared with the cells co-expressing mRFP-Q74 and control GFP **(Fig. 4c-e)**. Thus, our data indicates that chloroplast SPP can reduce the aggregation of polyQ proteins, which could have potential applications to ameliorate Huntington’s and other polyQ-based diseases.

**Figure 4.**
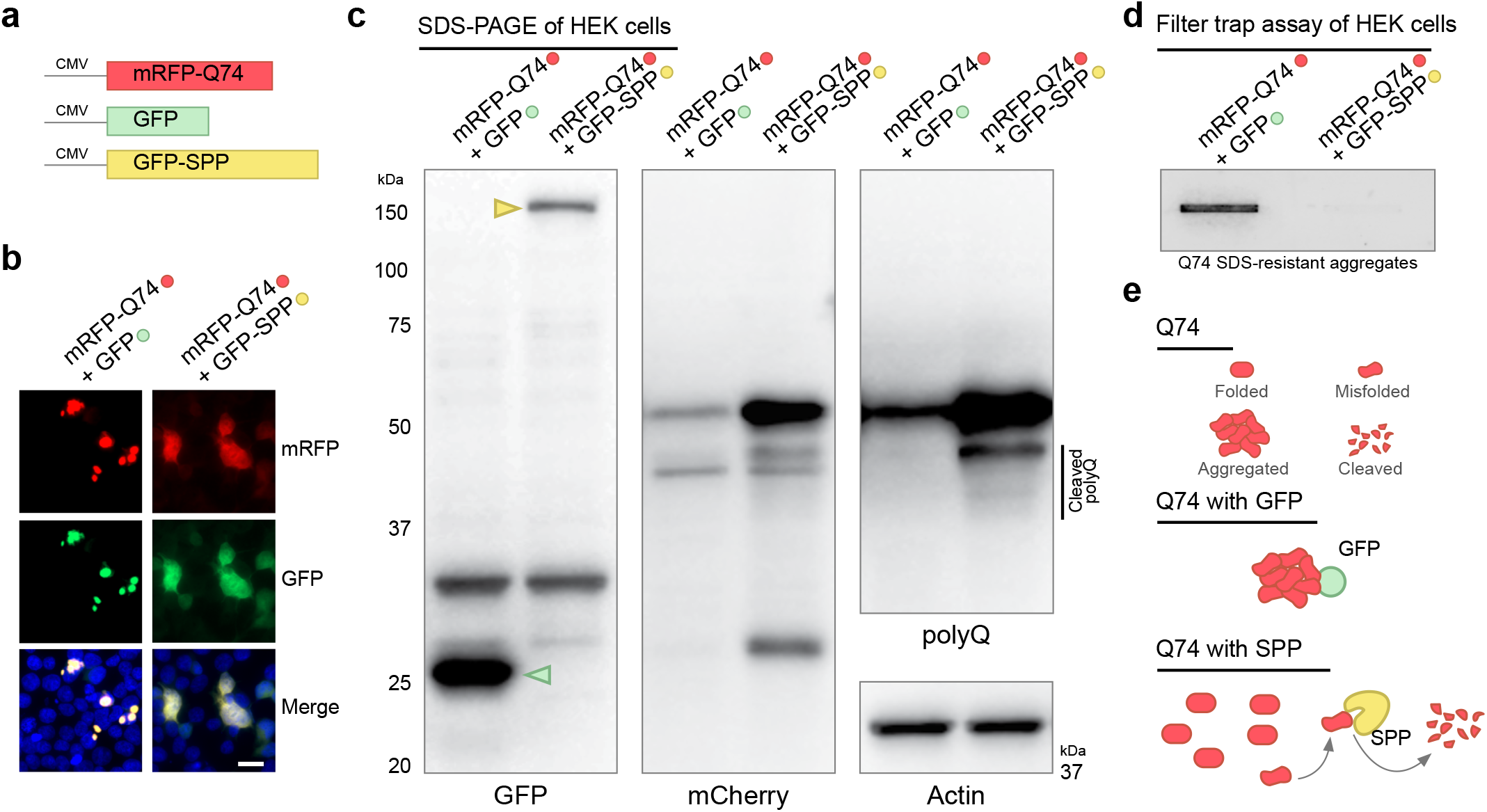
Synthetic SPP cleaves and reduces aggregation of pathogenic HTT exon1 Q74 in human cells. **(a)** Schematic representation of the constructs used for protein expression in HEK293 cells. (b) Microscopy analysis of HEK293 cells transfected with mRFP-Q74 and control GFP or mRFP-Q74 and GFP-SPP. Cell nuclei (in blue) were stained with Hoechst 33342. Scale indicates 20 μm. (c) SDS-PAGE of protein extracts of HEK293 cells transfected with mRFP-Q74 and control GFP or mRFP-Q74 and GFP-SPP. GFP antibody was used to detect free control GFP (27 kDa, indicated with a green arrowhead) and the human optimized SPP-GFP (approx. 163 kDa, indicated with a yellow arrowhead). mCherry and polyQ antibodies were used to detect complete and cleaved mRFP-Q74. Actin is shown as loading control. Representative images of 3 independent experiments are shown (d) Filter trap analysis showing mRFP-Q74 aggregation levels from samples indicated in c. (e) Schematic model of cleavage of Q74 by SPP. Misfolded version of Q74 are recognized and cleaved by SPP, avoiding the formation of aggregates.

### Chloroplast status coordinates the conformation and localization of the plastid Caseine Kinase 2 (pCK2)

Once we unveiled the connection between chloroplast proteostasis and regulation of Q69 aggregation, we asked whether LIN-treated chloroplasts can also promote the accumulation and/or aggregation of endogenous Arabidopsis proteins containing polyQ stretches **(Supplementary Data 1)**. To this end, we took advantage of a polyQ antibody which recognizes proteins containing polyQ stretches **(Supplementary Fig. 10)**. The treatment of wild-type Arabidopsis plants with LIN caused a strong accumulation and aggregation of polyQ-containing proteins **(Supplementary Fig. 10)**. Then, we focused on the plastid casein kinase 2 (pCK2), which is known to regulate chloroplast-to-nucleus communication (also referred to as retrograde signaling) by a still unknown mechanism (Wang et al. 2014; Rodiger et al. 2020). Remarkably, pCK2 contains a QQQQHQQQQQ region next to the chloroplast targeting sequence motif and prediction algorithms identified such polyQ region as a disorder and prion-like domain **(Fig 6a and Supplementary Fig. 9)**. In contrast to nuclear paralogs (nCK2s), the polyQ sequence of pCK2 and its prion-like domain is present in several plant species **(Supplementary Fig. 11-13)**, suggesting a conserved function for this polyQ region.

To examine a possible aggregation behavior of pCK2, we analyzed the distribution and localization of the full-length version of this protein fused to GFP after agroinfiltration of *N. benthamiana* leaves. To our surprise, we detected rounded GFP aggregates in the cytosol and GFP signal distributed homogenously in the nucleus and nucleolus **(Fig. 5b)**. Remarkably, chloroplast of plants agroinfiltrated with *35S:pCK2-GFP* showed rounded and amyloid-like fibrillar aggregates **(Fig. 5a-c)**. The amyloid-like structures formed a scaffold surrounding the chloroplast with a diamond-like structure **(Fig. 5c, d)**. Notably, the deletion of the polyQ domain reduced pCK2 aggregation, preventing the formation of the external chloroplast fibrillar scaffold **(Supplementary Fig. 14)**. We reasoned that LIN treatment could reduce import of pCK2 into the chloroplast and hence increase the formation of the fibrillar aggregates. To test this hypothesis, we infiltrated *N. benthamiana* plants expressing *35S:pCK2-GFP* with 800 μM LIN or mock solution. We found that the leaves infiltrated with LIN formed significantly more pCK2 diamond-shaped amyloid-like structures and have higher pCK2-GFP intensity in the nucleus compared to control mock-treated leaves **(Fig. 5d, e)**. Interestingly, infiltration with LIN did not increase pCK2ΔQ-GFP fluorescence in the nucleus, indicating that polyQ domain plays an important role in subcellular localization after chloroplast impairment. Together, our data indicate that chloroplast proteostasis status can regulate the conformation and localization of pCK2, providing a mechanistic explanation for its proposed role in retrograde signal **(Fig. 6)**.

**Figure 5.**
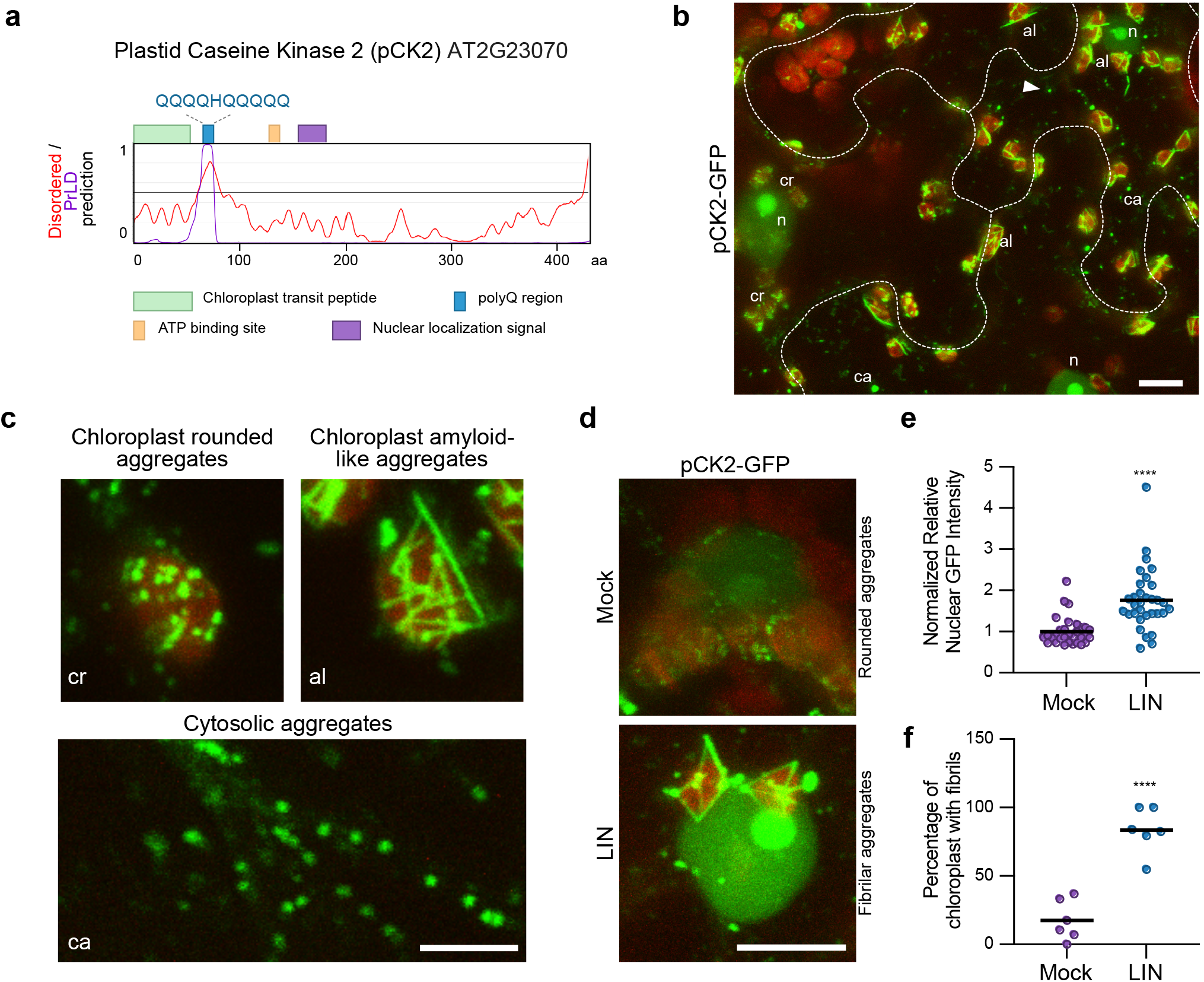
LIN treatment promotes fibrillar aggregation and nuclear localization of pCK2. **(a)** Schematic representation of the construct used for protein expression in *N. benthamiana* pavement cells. Scheme indicates position of different protein domains. Disordered (red line) and prion-like domain (purple line) prediction scores are also shown. **(b)** Confocal microscopy analysis of *N. benthamiana* pavement cells agroinfiltrated with *35S:pCK2-GFP*. Homogenously distributed GFP signal detected in nuclei (n). The image shows pCK2-GFP cytosolic aggregates (ca), chloroplast rounded aggregates (cr), and amyloid-like aggregates (al). Chloroplast autofluorescence is shown in red. Images were taken 3 dpa. Scale indicates 10 μm. **(c)** Closer magnification of chloroplast showing rounded aggregates (cr), chloroplast amyloid-like aggregates (al) and cytosolic aggregates (ca). Scale indicates 5 μm. **(d)** Representative confocal images of pCK2-GFP after 16 hours after infiltration of 800 μM LIN or mock solution. Leaves were infiltrated with LIN at day 3 after agroinfiltration. Scale indicates 10 μm. **(e)** Graph representing quantification of nuclear GFP intensity of LIN or mock treated cells expressing pCK2-GFP. Mean and individual values for mock (n = 31) and LIN (n = 32) are shown. (f) Graph representing the mean of the percentage of chloroplast with fibril structures after LIN and mock treatment. Chloroplast surrounding the nuclei from pavement cells were used for quantification in images taken at 40X. Data from six independent experiments. The statistical comparisons in e and f were made by two-tailed Student’s t test for unpaired samples. P value: ****P < 0.0001.

**Figure 6.**
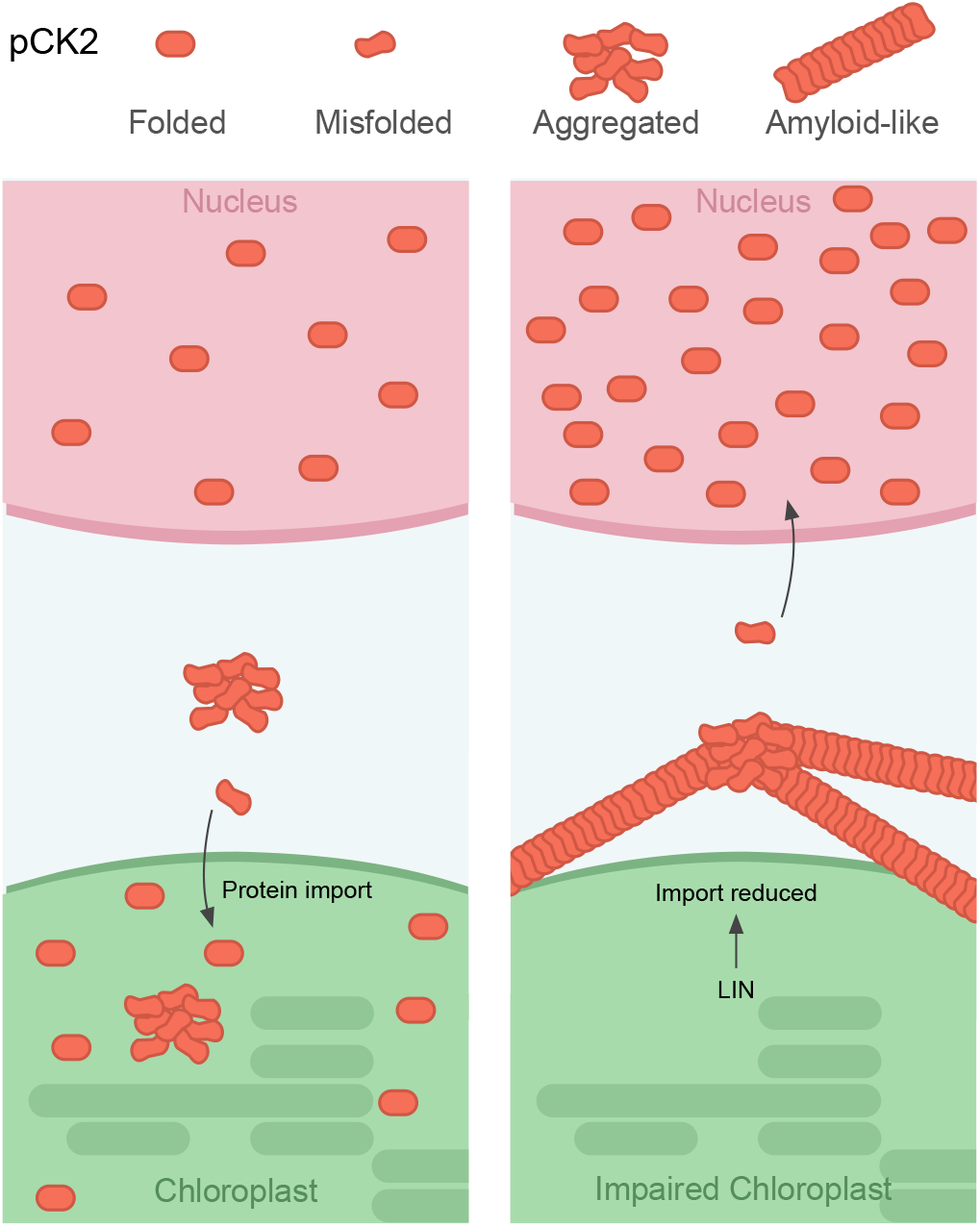
Model of pCK2 and its role in chloroplast-to-nucleus communication. Under normal conditions (left panel), pCK2 accumulates, in nucleus cytosol, and chloroplast. However, when chloroplast protein import is reduced using LIN (right panel), pCK2 accumulates surrounding the chloroplast in amyloid-like fibrils and relocates to higher levels to the nucleus. Similar to nCK2s, pCK2 in the nucleus could phosphorylate several targets modulating gene expression.

## Discussion

### Chloroplasts are main posttranslational regulators of polyQ-extended proteins

Plants are known for their striking resilience to abiotic stress due to their enhanced components of the proteostasis network (Finka et al. 2011; Llamas et al. 2021). To our knowledge, in contrast to mammals, plants do not suffer from proteinopathies or conformational disorders caused by structural abnormal proteins like amyloids. In plants, amyloid formation mediates protein storage in seeds (Antonets et al. 2020). To better understand the enhanced proteostasis network of plants, we challenged *Arabidopsis thaliana* by expressing the toxic fragment of the human mutant HTT, which is the pathological base of Huntington’s disease.

Expression of the pathogenic fragment of the exon 1 HTT with more than 35Q repeats cause its aggregation, disrupting cellular functions and eventually leading to cell death in several model organisms such as yeast, nematodes, and human cells (Pearce and Kopito 2018; Yang and Yang 2020; Peskett et al. 2018). However, the constitutive expression of exon 1 HTT with 69 Q repetitions (here referred to as Q69) did not cause toxic or deleterious effects in Arabidopsis plants **(Fig. 1d-f and Supplementary Fig. 2)**. The interactome of Q28 and Q69 in Arabidopsis showed that both proteins interact with several protein orthologues that were previously identified as polyQ regulators (Wang et al. 2006; Kim et al. 2016). Unexpectedly, polyQ proteins in plants also interact with components of the chloroplast proteostasis network **(Fig. 2)**. Compared to other eukaryotic organisms, the presence of chloroplasts in plant cells has expanded the variety of proteostasis components such as chaperones and proteases that may also counteract cytosolic toxic protein aggregation. In non-plant models, it has been proposed that aggregated cytosolic proteins are disentangled on the mitochondria surface and imported to enable their degradation by mitochondrial proteases (Ruan et al. 2017; Li et al. 2019; Schlagowski et al. 2021). Our proteomics data of polyQ interacting proteins in Arabidopsis provided a first indication that plant chloroplasts can import and degrade cytosolic misfolded polyQ proteins.

We found that impairing chloroplast import, either pharmacologically or genetically, trigger the formation of Q69 aggregates **(Fig. 3)**. The unstructured configuration of Q69 led us to hypothesize that the polyQ region could be recognized as an unfolded N-terminal transit peptide in a pre-protein. In support of this idea, we found that Q69 interacts with SPP which recognizes, binds, and cleaves the transit peptides of chloroplast pre-proteins (Zhong et al. 2003). Using human cell lines, we demonstrated that a synthetic SPP can cleave and reduce aggregation of extended polyQ proteins. Together, our data strongly suggest that misfolded versions of Q69 are recognized by the chloroplast import machinery for further degradation by SPP and possibly by Clp and FtsH proteases. Importantly, the cleavage of polyQ expanded proteins by SPP in human cells can open the door to novel plant-based strategies to ameliorate polyQ-caused disorders such as Huntington’s and Machado-Joseph diseases. localization of pCK2 to the nucleus was dependent on the polyQ stretch **(Supplementary Fig. 14)**, indicating that pCK2 prion-like domain is important for retrograde signaling. A recent study confirmed, for the first time in plants, the formation of amyloids in the accumulation of storage proteins in seeds (Antonets et al. 2020). However, the functional and physiological relevance of pCK2 amyloid-like structures remains to be elucidated. We speculate that pCK2 amyloid-like structures may work as stress molecular memory device dependent on its conformational changes, based on the suggested role prion-like proteins in plants (Garai et al. 2021; Jung et al. 2020; Dorone et al. 2021).

### A polyQ-prion-like domain in the plastid casein kinase 2 controls its conformation and localization

The results suggesting that Q69 is imported and degraded in chloroplasts led us to hypothesize that such organelles could also regulate the proteostasis status of intrinsic Arabidopsis proteins with polyQ stretches **(Supplementary Data 1)**. Indeed, filter trap assays using a polyQ antibody indicated that several Arabidopsis proteins aggregate after short LIN treatment **(Supplementary Fig. 10)**. Interestingly, among our polyQ-containing proteins with a predicted chloroplast transit peptide **(Supplementary Fig. 8)**, we found the plastid casein kinase (pCK2) as an interesting candidate to characterize its aggregation propensity. pCK2 harbors a chloroplast transit peptide and a nuclear localization signal **(Fig 5a)** (Salinas et al. 2006). pCK2 has been termed plastid transcription kinase because of its localization and activity on the plastid transcription system (Baginsky et al. 1997; Rodiger et al. 2020). Arabidopsis contains four CK2 alpha subunits, three localized in nucleus and cytoplasm and pCK2 which was reported to be exclusively located in chloroplasts (Salinas et al. 2006; Mulekar and Huq 2015). Our results indicate that pCK2, like the other CK2 alpha subunits, can also localize in cytosol and nucleus **(Fig. 5b-d)**. Interestingly, treatment with LIN promotes the formation of amyloid-like fibrillar structures encircling the chloroplast and increases the localization of pCK2 in the nucleus **(Fig. 5c-e)**. These results could explain how pCK2 plays a dominant role over nuclear CK2s in regulating gene expression during abscisic acid (ABA) and heat-stress signaling (Wang et al. 2014). Notably, mutant *pCK2* plants display reduced expression of genes involved in retrograde signaling such as *ABI4, Prs1* and *Hsps* (Wang et al. 2014). The dual localization nature of pCK2 could regulate gene expression in both nucleus and chloroplast.

The formation of fibrillar aggregates and increased localization of pCK2 to the nucleus was dependent on the polyQ stretch **(Supplementary Fig. 14)**, indicating that pCK2 prion-like domain is important for retrograde signaling. A recent study confirmed, for the first time in plants, the formation of amyloids in the accumulation of storage proteins in seeds (Antonets et al. 2020). However, the functional and physiological relevance of pCK2 amyloid-like structures remains to be elucidated. We speculate that pCK2 amyloid-like structures may work as stress molecular memory device dependent on its conformational changes, based on the suggested role prion-like proteins in plants (Garai et al. 2021; Jung et al. 2020; Dorone et al. 2021).

### Retrograde signaling pathways involve dual-localized proteins

The mechanism discovered here, whereby the chloroplast status directly influences the subcellular distribution of pCK2 and consequently the access to its biological targets, offers a new framework for a role of this protein in chloroplast-to-nucleus communication **(Fig. 6)**. The proposed mechanism resembles that of the mitochondrial unfolded protein response (mtUPR) in *Caenorhabditis elegans*. When protein import is impaired, the bZIP transcription factor ATFS-1 translocates to the nucleus and activates the mtUPR. Notably, ATFS-1 harbors a nuclear localization signal and a mitochondrial transit peptide, and the relatively weak transit peptide allows the transcription factor to serve as a sensor of mitochondrial import efficiency (Nargund et al. 2012; Melber and Haynes 2018). Using the predictor software PLAAC, we found that ATFS-1 has a prion-like domain rich in Qs next to the transit peptide, similar to the location of the polyQ domain in pCK2. Our data suggest that the presence of nonpathological prion-like domains may be a conserved mechanism to regulate organelle import and trafficking impacting on retrograde signaling and gene expression. Like ATFS-1, the Arabidopsis transcription factor WHIRLY1 (WHY1) relocate form chloroplast to nucleus upon changes in the redox state of the chloroplast (Isemer et al. 2012; Foyer et al. 2014). We demonstrated that several proteins containing polyQ domains aggregate shortly after LIN treatment **(Supplementary Fig. 10)**. Interestingly, many of the polyQ-containing proteins in Arabidopsis proteome are annotated as transcription factors **(Supplementary Data 1)**. Based on our data, we speculate that the status of the chloroplast could change the conformation and activity of several transcription factors hence having an important role on nuclear gene expression.

### Terminating misfolded/aggregated proteins to target aging and climate change

Unpredictable daily temperature fluctuations due to global climate change trigger protein misfolding and aggregation resulting in decreased plant growth and productivity (Llamas et al. 2021; Zhao et al. 2017). Modulating retrograde signaling pathways and rewiring proteostasis mechanisms in crop plants could help to maintain plant productivity even in the face of frequent temperature changes. Overexpression or fine modulation of some of the identified polyQ-interactor proteins could be used as a strategy to generate stress-resistant crops with reduced levels of stress-generated protein aggregates.

A hallmark event in age-related and neurodegenerative disease is the accumulation of misfolded/aggregated proteins that lead to cellular dysfunction and cell death (Soto and Pritzkow 2018). The mechanism described here whereby a chloroplast synthetic SPP decreases expanded-polyQ aggregation in human cells, opens a possibility to develop novel treatments against human polyQ-based neurodegenerative diseases. However, further work is required to unveil the detailed molecular mechanisms by which SPP binds and cleaves misfolded polyQ proteins. In these lines, it will be fascinating to explore whether the synthetic SPP, or other plant proteins, can also be used to clear toxic aggregates of TDP-43 and FUS which cause amyotrophic lateral sclerosis (ALS) (Hommen et al. 2021).

## Materials and methods

### Plant material, constructs, and growth conditions

All the Arabidopsis thaliana lines used in this work are in Columbia-0 ecotype. WT, and *toc159* (Woodson et al. 2015; Ling et al. 2019) were used in this study. All seeds were surface-sterilized and germinated on solid 0.5X Murashige and Skoog (MS) medium with vitamins without sucrose. Plants were incubated in a growth chamber at 22 ºC (or otherwise indicated) under long-day conditions (or otherwise indicated). When indicated medium was supplemented with MG-132 (bio-techne) or LIN (Sigma). LIN was applied in *N. benthamiana* leaves by direct infiltration with a plastic syringe. Liquid 0.5X MS medium with vitamins without sucrose were supplemented with LIN or MG-132. For the root growth analysis, we used FIJI (ImageJ) to measure the root length from 7-day-old seedlings grew in vertical agar plates. BP Clonase II Enzyme mix (Thermo Fisher) and Gateway LR Clonase II Enzyme mix (Thermo Fisher) were used for cloning. *Q28* and *Q69* genes were obtained by PCR using as a template the plasmid pEGFP-Q74. Different polyQ length products were amplified and sequenced. *Q28* and *Q69* genes were subcloned in the entry vector pDONR221 and then cloned in the vector pMpGWB105 (Q constructs). The full-length coding sequence of Arabidopsis *pCK2* (AT2G23070) was amplified by PCR and the synthetic *pCK2ΔQ* was ordered (IDT). *pCK2* and *pCK2ΔQ* were cloned in the pGWB505 vector. Cultures of *Agrobacterium tumefaciens* GV3101 containing constructs of interest were agroinfiltrated in *N. benthamiana* leaves as previously described (Perello et al. 2016). Arabidopsis transgenic plants were generated by the floral dip method (Clough and Bent 1998). The *35S:Citrine-Q69* transgene was introgressed into the *toc159* mutant background by cross-fertilization. For the flowering time experiment, 30 plants from each line were germinated and grown in short-day conditions, rosette leaves number were counted until the production of a visible bolt. For the heat shock assays, single plate containing 7-day-old WT, Q28 and Q69 was covered with aluminum foil and transferred to 45 °C (or 37 ºC) for indicated times. Mock plate was also cover with aluminum foil and left at control conditions. The heat-treated plates were transferred back to light conditions at 22°C.

### Microscopy and fluorescence quantification

Confocal microscopy images were taken with a Meta 710 Confocal Microscope with laser ablation 266 nm (Zeiss). All images were acquired using the same parameters between experiments. For imaging human HEK cells Imager Z1 microscope (Zeiss) was used. Quantification of GFP fluorescence was performed with FIJI (ImageJ) software.

### Gene expression analysis

Total RNA was extracted from plant tissues, using the RNeasy Plant Mini Kit (Qiagen). RNA was quantified using a NanoDrop (Thermo Scientific). cDNA was synthetized using the qScript Flex cDNA synthesis kit (Quantabio). SybrGreen real-time quantitative PCR experiments were performed with a 1:20 dilution of cDNA using a CFC384 Real-Time System (Bio-Rad). Data were analyzed with the comparative 2ΔΔCt method using the geometric mean of *Ef1α* (5’-CTG GAG GTT TTG AGG CTG GTA T-3’ and 5’-CCA AGG GTG AAA GCA AGA AGA-3’) and *PP2A* (5’-TAA CGT GGC CAA AAT GAT GC-3’ and 5’-GTT CTC CAC AAC CGC TTG GT-3’) as housekeeping genes. Following primers were used: *Hsc70-1* (5’-AAG GAA ACA GAA CCA CGC CA-3’ and 5’-TGT CAG AGA AAC GAC GAC CG-3’), *Hsp70b* (5’-GCA GAA GAT TGA GAA GGC GAT TG-3’ and 5’-CGC TCT AAT CCA CCT CTT CGA TC-3’), *Hsp90-1* (5’-AAG CTC GAT GGA CAG CCT GAA C-3’ and 5’-TCC CAA GTT GTT CAC CAA ATC TGC-3’), and *Hsp101b* (5’-GAG GAG TTG CTT TGG CAG TC-3’ and 5’-CAG CGC CTG CAT CTA TGT AA-3’).

### Statistical analysis

PRISM 9 software was used for statistical analysis.

### Analysis of the Arabidopsis polyQ proteome

The whole review and manually annotated Arabidopsis proteome was downloaded from UniProt. Protein sequences were filtered when containing more than ≥ 5 consecutive Q repeats. Annotated chloroplast proteins were also filtered. Selected protein sequences were analyzed for the presence of prion like domains (PrLDs) using the PLAAC software (http://plaac.wi.mit.edu/details) (Lancaster et al. 2014). A minimum length for prion domains (L core) was set at 60 and parameter α was set at 50. For background frequencies, A. thaliana proteome was selected. To identify intrinsically disordered regions (IDRs) we used the IUPred3 software (https://iupred.elte.hu/) (Erdos et al. 2021). We performed the IUPred3 long disorder analysis with medium smoothing.

### Filter trap analyses

Protein extracts were obtained with native lysis buffer (300 mM NaCl, 100 mM Hepes pH 7.4, 2 mM EDTA, 2% Triton X-100) supplemented with 1X plant protease inhibitor (Merck). Cellular debris was removed by several centrifugation steps at 8,000 x g for 10 min at 4 °C. Supernatant was recollected and protein concentration determined with the Pierce BCA Protein Assay Kit (Thermo Fisher). A cellulose acetate membrane filter (GE Healthcare Life Sciences) was placed in a slot blot apparatus (Bio-Rad) coupled to a vacuum system. Membrane was equilibrated with 3 washes with equilibration buffer (native buffer supplemented with 0.5% SDS). 300, 200, and 100 µg of protein extract was supplemented with SDS at a final concentration of 0.5% and loaded and filter through the membrane. Then, the membrane was washed with 0.2% SDS. The membrane was blocked in 3% BSA in TBST for 30 min followed by 3 washes with TBST. Membrane was incubated either with anti-GFP [1:2,000] (Abcam, ab6556) or anti-mCherry [1:5,000] (Abcam, ab167453). Membrane was washed 3 times for 5 min and incubate with secondary antibodies in TBST 3% BSA for 30 min. Membrane was developed using an Odissey DLx (Licor). Extracts were also analyzed by SDS-PAGE and western blotting to determine loading controls.

### Western blot analysis

Plant material was recollected and grinded in liquid N2. The powder was resuspended on ice-cold TKMES homogenization buffer (100 mM Tricine-potassium hydroxide pH 7.5, 10 mM KCl, 1 mM MgCl_2_, 1 mM EDTA, and 10% [w/v] Sucrose) supplemented with 0.2% (v/v) Triton X-100, 1mM DTT, 100 µg/ml PMSF, 3 µg/ml E64, and 1X plant protease inhibitor (Sigma). The resuspended sample was centrifuged at 10,000 × g for 10 min at 4 °C and the supernatant recovered for a second step of centrifugation. Protein concentration was determined using the kit Pierce Coomassie Plus (Bradford) Protein-Assay (Thermo Scientific). Total protein was separated by SDS–PAGE, transferred to nitrocellulose membrane, and subjected to immunoblotting. The following antibodies were used anti-GFP [1:2,000] (Abcam, ab6556), anti-mCherry [1:5,000] (Abcam, ab167453), anti-Actin [1:5000] (Agrisera, AS132640), anti-Actin [1:5000] (Abcam, ab8226), anti-polyQ [1:1,000] (Merck, MAB1574), anti-Hsp90-1 (Agrisera, AS08346), anti-Hsp70 (Agrisera, AS08371), and anti-ATG8 (Agrisera, AS142769).

Membranes were visualized by chemiluminescence. *C. elegans* strain AM23 (rmIs298[F25B3.3p::Q19::CFP]) (Brignull et al. 2006) were lysed with native lysis buffer and used for immunoblotting.

### Experiments with HEK293T cells

*CMV:pEGFP-Q74* plasmid was digested with BglII and BamHI to remove *Q74* gene and generate *pEGFP* (*CMV:GFP*). The *SPP* (AT5G42390) gene, codon optimized and without chloroplast transit peptide was synthetized (Twist Bioscience). To generate *CMV:GFP-SPP*, the synthetic gene was cloned in the *pDEST-CMV-N-GFP* vector by Gateway technology. *CMV:mRFP-Q74, CMV:GFP-SPP*, and *CMV:GFP* were used for transfection. HEK293T cells (ATCC) were plated on 0.1% gelatin-coated plates and grown in DMEM (Gibco) supplemented with 10% fetal bovine serum and 1% MEM non-essential amino acids (Gibco) at 37 °C, 5% CO2 conditions. HEK293T cells were transfected with the constructs approximately 24 h after seeding with 1 μg of each construct. DNA was incubated at 80°C for 5 min and mixed with FuGENE HD (Promega) in a ratio of 3:1 (FuGENE:DNA) and 65 μl of Opti-MEM (Thermo Fisher) was added. The mix was incubated at room temperature for 15 min. The total mix was added dropwise to the cells. HEK293 cells were incubated at 37°C and 5% CO_2_, refreshing every day with supplemented DMEM. After 72 h of incubation, cells were harvested for further experiments. For microscopy analysis cells grown on coverslips were washed two times with 1 ml of 1x DPBS (Gibco). Then, cells were fixed with 1 ml of 4% paraformaldehyde (PFA) solution for 20 min, followed by 2 washing steps with 1 ml of DPBS. Cells were mounted in mounting solution, on microscope slides for further microscopy analysis. For filter trap and western blot, cells were lysed in non-denaturing native lysis buffer, scraped from the tissue culture plates, and homogenized through a syringe needle (27G). Samples were centrifuged at 10,000 x g for 10 min at 4 °C and supernatant was collected. Protein concentration was determined with the Pierce BCA Protein Assay Kit (Thermo Fisher).

### Phylogenetic analysis

For the identification of CK2 proteins in photosynthetic organisms systematic BLASTP analyses (blast.ncbi.nlm.nih.gov/Blast.cgi and phytozome-next.jgi.doe. gov/blast-search) were performed using the full-length Arabidopsis pCK2 (AT2G23070), nCK2A (AT5G67380), nCK2B (AT3G50000), and nCK2C (AT2G23080) proteins. Sequence alignments were performed using CLUSTALW (www.ebi.ac.uk/Tools/msa/clustalo). Subsequently, phylogenetic trees were constructed based on the complete protein sequences. The trees were constructed by rooting at midpoint using the maximum likelihood methods in MEGA11 program (http://www.megasoftware.net) (Tamura et al. 2021). For estimating the reliability of the trees, we performed bootstrapping using 2000 replications (Hall 2013). The evolutionary distances were computed using the Jones-Taylor-Thornton correction model.

### PAM fluorometric measurements

For the assessment of the photosynthetic activity, the treated plant samples were incubated for 20 min in total darkness to reach a dark-adapted state. The photosynthetic activity of the plants was measured using the M-Series PAM fluorometer (Heinz Walz GmbH, Effeltrich, Germany). The data was evaluated using the ImagingWin software (v.2.59p; Walz, Germany).

### Proteomics analysis

For label-free quantitative proteomics, 7-day-old Q28 and Q69 seedlings were used. Seedlings were lysed in lysis buffer (1% Triton X-100, 50 mM Tris–HCl pH 8.0) supplemented with 1X plant protease inhibitor cocktail (Sigma) and 25 mM *N*-ethylmaleimide. Samples were homogenized by vortexing and centrifuged at 13,000 x g for 10 min at 4 °C. Then the protein lysates were incubated for 1 hour with anti-GFP (ImmunoKontakt (Amsbio), TP401, 1:500). As a co-immunoprecipitation control, the same amount of protein lysates was incubated with IgG (Abcam, ab46540) in parallel. After antibody incubation, samples were incubated with 50 μl of μMACS Micro Beads for 1 hour at 4 °C with overhead shaking. Then, samples were loaded to pre-cleared μMACS column (#130-042-701). After loading samples, beads were washed three times with washing buffer 1 (50 mM Tris (pH 7.5) buffer containing 150 mM NaCl, 5% glycerol and 0.05% Triton) and then washed five times with washing buffer 2 (50 mM Tris (pH 7.5) and 150 mM NaCl). Then, in-column trypsin digestion was performed with trypsin digestion solution (7.5 mM ammonium bicarbonate, 2 M urea, 1 mM DTT and 5 ng ml^−1^ trypsin). Trypsinazed samples were eluted with two times 50 μl of elution buffer (2 M urea, 7.5 mM Ambic, and 10 mM chloroacetamide). Then, samples were incubated overnight at room temperature with shaking in the dark. Samples were stage-tipped the next day for label-free quantitative proteomics. All samples were analyzed on a Q-Exactive Plus (Thermo Scientific) mass spectrometer coupled to an EASY nLC 1200 UPLC (Thermo Scientific). Peptides were loaded with solvent A (0.1% formic acid in water) onto an inhouse packed analytical column (50 cm × 75 μm I.D., filled with 2.7 μm Poroshell EC120 C18 (Agilent)). Peptides were chromatographically separated at a constant flow rate of 250 nl min^−1^ using the 150-min method: 3–5% solvent B (0.1% formic acid in 80% acetonitrile) within 1 min, 5–30% solvent B (0.1% formic acid in 80% acetonitrile) within 65 min, 30–50% solvent B within 13 min and 50–95% solvent B within 1 min, followed by washing and column equilibration. The mass spectrometer was operated in data-dependent acquisition mode. The MS1 survey scan was acquired from 300 to 1,750 *m/z* at a resolution of 70,000. The top 10 most abundant peptides were subjected to higher collisional dissociation fragmentation at a normalized collision energy of 27% and the automatic gain control target was set to 5 × 10^5^ charges. Product ions were detected in the Orbitrap at a resolution of 17,500. All mass spectrometric raw data were processed with MaxQuant (version 1.5.3.8) using default parameters as described above. LFQ was performed using the LFQ mode and MaxQuant default settings. All downstream analyses were carried out on LFQ values with Perseus (version 1.6.2.3). MS2 spectra were searched against the *Arabidopsis thaliana* Uniprot database, including a list of common contaminants. False discovery rates (FDRs) on protein and peptide–spectrum match (PSM) level were estimated by the target-decoy approach to 0.01% (Protein FDR) and 0.01% (PSM FDR) respectively. The minimal peptide length was set to 7 amino acids and carbamidomethylation at cysteine residues was considered as a fixed modification. Oxidation (M) and Acetyl (Protein N-term) were included as variable modifications. The match-between runs option was enabled. Label-free quantification (LFQ) was enabled using default settings. The resulting output was processed using Perseus as follows: protein groups flagged as “reverse”, “potential contaminant” or “only identified by site” were removed from the proteinGroups. txt. LFQ values were log_2_ transformed. Missing values were replaced using an imputation-based approach (random sampling from a normal distribution using a down shift of 1.8 and a width of 0.3). Significant differences between the groups were assessed using Student’s t-test. A permutation-based FDR approach was applied to correct for multiple testing. Data visualization was done with Instant Clue.

### Immunoprecipitation

7-day-old Q28 and Q69 seedlings were lysed in lysis buffer (1% Triton X-100, 50 mM Tris–HCl pH 8.0) supplemented with protease inhibitor cocktail (Roche) and 25 mM N-ethylmaleimide. Samples were homogenized by vortexing and centrifuged at 13,000 x g for 10 minutes at 4 °C. Then the protein lysates were incubated for 1 hour with anti-GFP (ImmunoKontakt (Amsbio), TP401, 1:500). As a co-immunoprecipitation control, the same amount of protein lysates was incubated with IgG (Abcam, ab46540) in parallel. After antibody incubation, samples were incubated with 50 μl of μMACS Micro Beads for 1 h at 4 °C with overhead shaking. Then, samples were loaded to pre-cleared μMACS column (#130-042-701). After loading samples, beads were washed three times with washing buffer 1 (50 mM Tris (pH 7.5) buffer containing 150 mM NaCl, 5% glycerol and 0.05% Triton X-100) and then washed five times with washing buffer 2 (50 mM Tris (pH 7.5) and 150 mM NaCl). The beads were subjected to 1× Laemmli buffer for 5 min for elution and collection. The samples were boiled for 5 min at 95 °C and were used for SDS–PAGE and western blot analysis.

## Contributions

E.L. conceptualized the project, designed and performed most of the experiments, analyzed the data, and wrote the manuscript. S.K. performed immunoprecipitation experiments and R.G.G. analyzed proteomics data. H.J.L performed experiments with human cells. N.D., N.C., S.T.M., and E.S. helped to perform some of the experiments. P.P. performed the phylogenetic analysis of pCK2. M.R.C. and A.Z. interpreted and discussed the results and provided reagents and equipment for the research. D.V. planned and supervised the project and wrote the manuscript. All authors discussed the results and commented on the manuscript.

## Acknowledgments

This work was supported by the European Research Council (ERC Starting Grant-677427 StemProteostasis), the Deutsche Forschungsgemeinschaft (DFG) (Germany’s Excellence Strategy-CECAD, EXC 2030-390661388 / Germany’s Excellence Strategy – EXC-2048/1 – project ID 390686111 (CEPLAS) / SFB-1403–414786233). E. Llamas was supported with a Humboldt postdoctoral fellowship. We thank T. Kohchi for the pMp-GWB105 (Addgene 68559), T. Nakagawa for the pGWB502 (Addgene 74844), D. Rubinsztein for the pEGFP-Q74 (Addgene 40262) and R. Ketteler for the pDEST-CMV-N-EGFP (Addgene 122842). We thank R.I. Morimoto for providing the C. elegans strain AM23 (rmIs298[F25B3.3p::Q19::CFP]). P. Pulido thanks the Community of Madrid (grant 2019-T1/BIO-13731) and the Spanish Agencia Estatal de Investigación (grant PID2020118607RB-I00). We thank CECAD (A. Schauss and C. Jüngst) imaging facilities for their support in confocal microscopy. We thank the CECAD Proteomics Facility for contribution and advice on proteomics experiments.

