## Supplementary Figures for "Chloroplast protein import determines plant proteostasis and retrograde signaling"

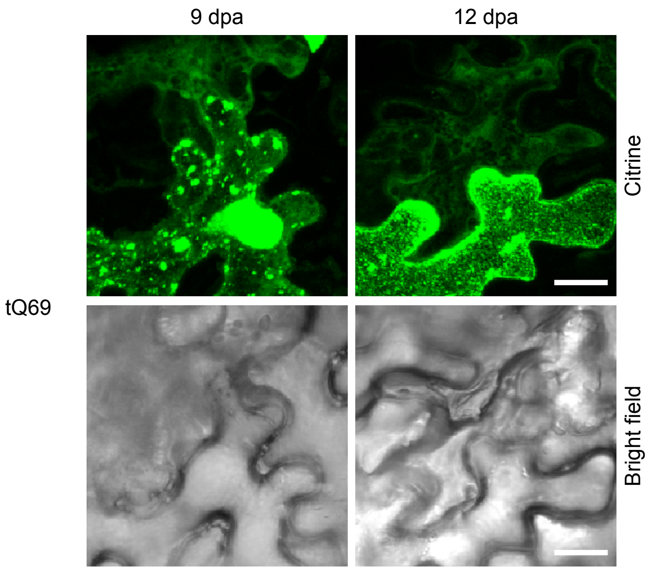


**Supplementary Figure 1. Heterogenous aggregation of tQ69 in epidermal pavement cells.** Images show representative epidermal cells transiently expressing tQ69 at 9 and 12 days post-agroinfiltration (dpa) from 4-weeks-old *Nicotiana benthamiana* agroinfiltrated leaves. Citrine and Bright field are shown. Scale indicates 20 µm.

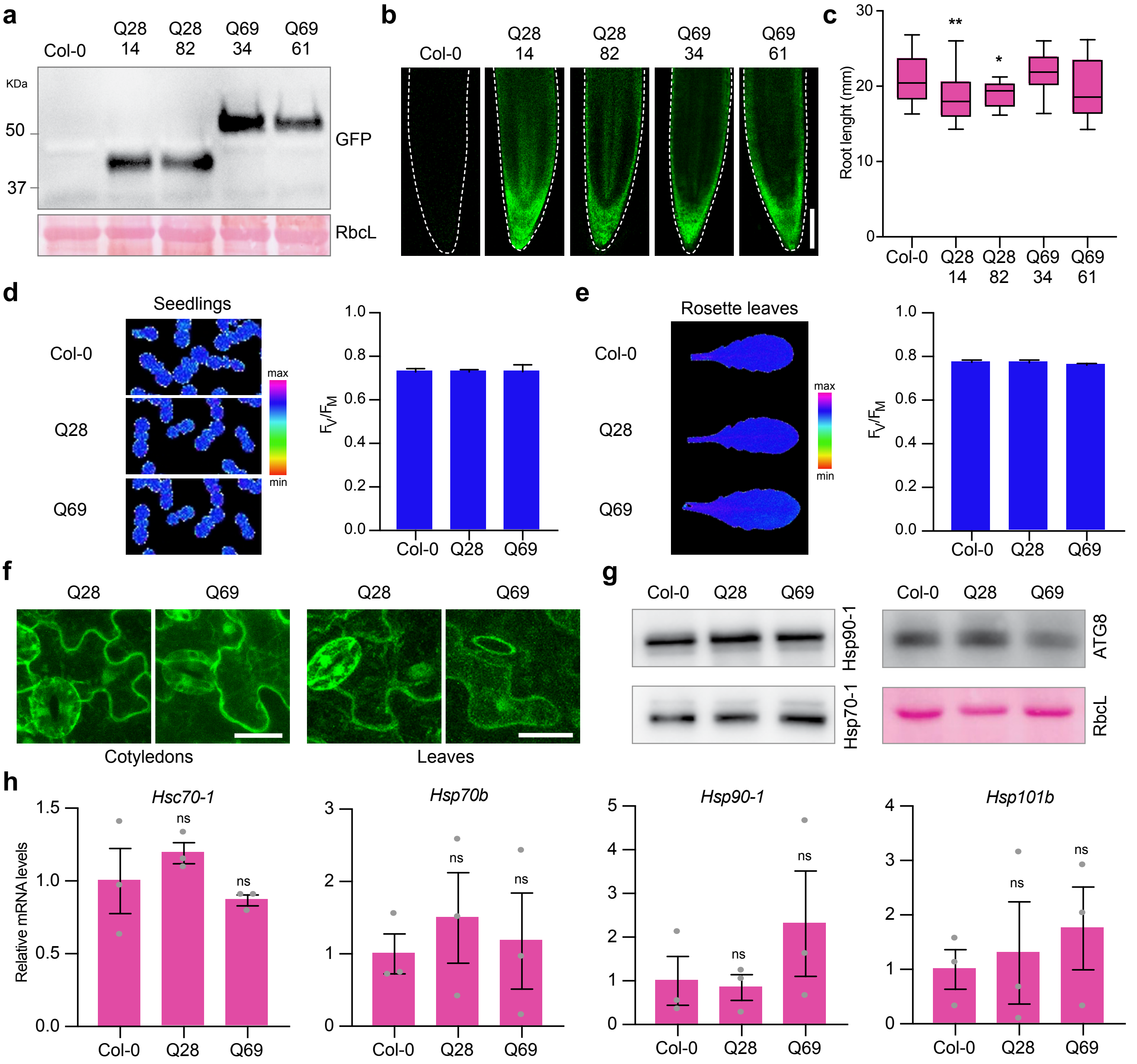


**Supplementary Figure 2. Characterization of constitutive Q28 and Q69 plants. (a)** Immunoblot analysis against GFP of independent transgenic Q28 and Q69 plants. Ponceau S staining showing RbcL was used as loading control. **(b)** Microscopy analysis of 7-day-old Arabidopsis transgenic plants expressing Q28 and Q69. Scale indicates 100 µm. **(c)** Root growth analysis of 7-day-old Col-0, Q28 and Q69 seedlings (n = 30 roots). **(d)** Photosynthetic activity activity $F_V/F_M$ of 7-day-old Col-0, Q28 and Q69 seedlings and **(e)** of rosette leaves of 38-day-old plants measured via PAM fluorometry. Data from two independent measurements. **(f)** Confocal microscopy analysis showing the distribution of Q28 and Q69 in cotyledons of 7-day-old seedlings and the fourth true rosette leaves of 22-day-old Arabidopsis plants. Scales indicate 20 µm. **(g)** Immunoblot analysis of proteostasis marker proteins. Samples of 7-day-old seedlings were used. RbcL was used as loading control. **(h)** Proteostasis-related genes are not upregulated in polyQ lines compared to Col-0. 7-day-old-seedlings grown under normal conditions (22 ºC) were used. The statistical comparisons in c and h were made by two-tailed Student’s t-test for unpaired samples. P value: *P < 0.05, **P < 0.01, ns = not significant (P > 0.05).

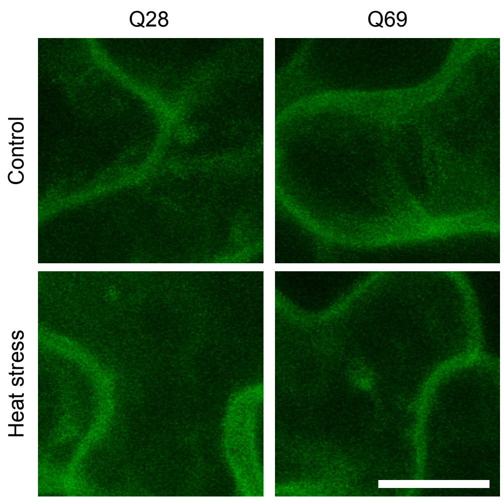


**Supplementary Figure 3. 37 ºC heat shock treatment does not cause polyQ aggregation.** (a) Images show representative epidermal cells from cotyledons. 7-day-old seedlings grown at 22 ºC were transfer in dark to incubators at 37 ºC (Heat stress) or 22 ºC (control) for 90 minutes. Scale indicates 10 μm.

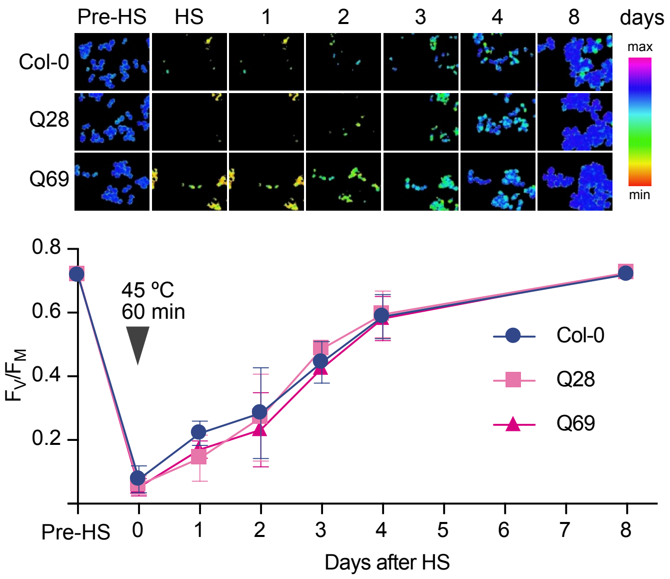


**Supplementary Figure 4. PolyQ plants do not show increased sensitivity to heat stress compared to WT.** Col-0, Q28 and Q69 7-day-old-plants grown at 22 ºC in short-day (SD) conditions were transfer to 45 ºC for 60 minutes. After heat shock (HS) plant were transfer back to 22 ºC (SD). Photosynthetic parameters were measured before HS, immediately after HS, and at day 1, 2, 3, 4, and 8 after HS. Upper panel shows the visualization of $F_V/F_M$ measured via PAM fluorometry. Blue indicates a high photosynthetic activity while green decreased and yellow strongly decreased photosynthetic activity. Graph represents the mean ± s.e.m of 4 independent experiments. The statistical comparisons were made by two-tailed Student's t-test for un-paired samples were not significant (P > 0.05) differences were found.

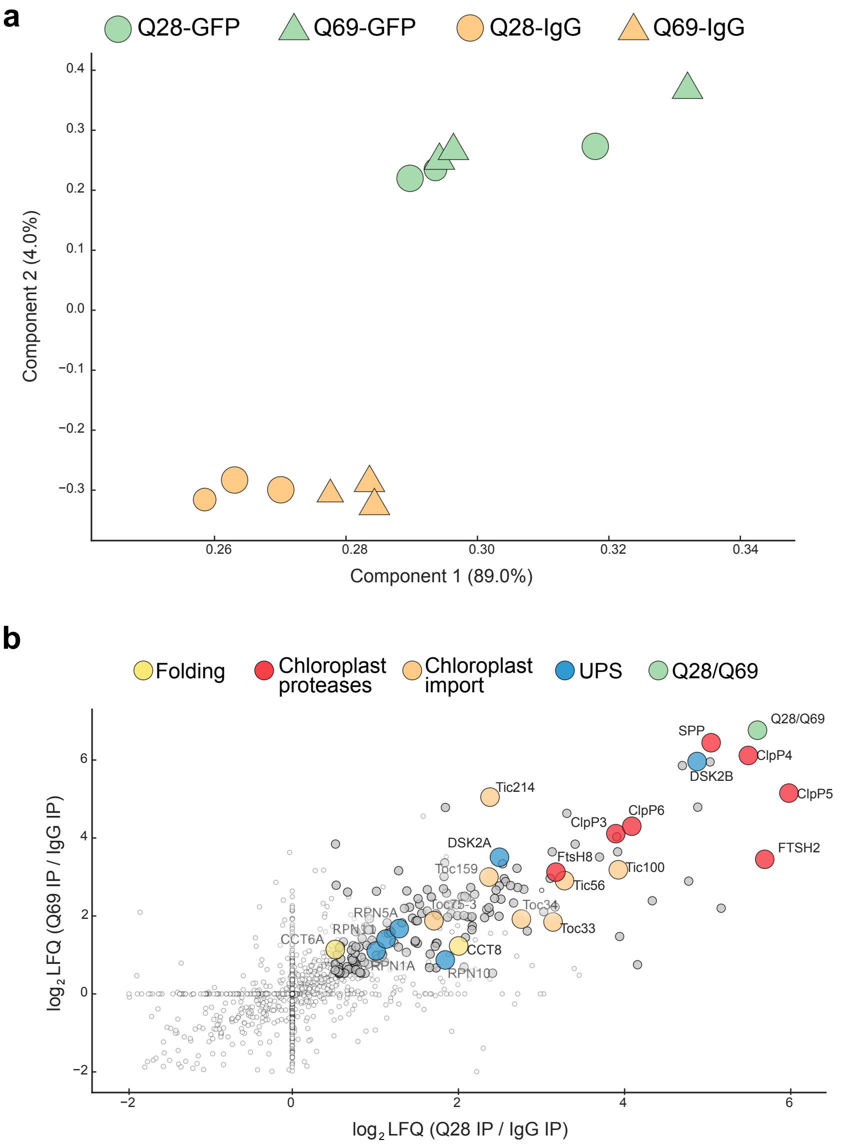


**Supplementary Figure 5. Q28 and Q69 share common protein interactors. (a)** Principal component analysis (PCA) groups Q28 and Q69 interactors (n = 3). **(b)** Scatterplot of protein enrichments in Q28/Q69 co-IP from Q28 and Q69 7-day-old seedlings. Gray and colored circles indicate significance after correction for multiple testing (false discovery rate (FDR) < 0.05 was considered significant). Yellow colored circles represent proteins involved in protein folding, red colored proteins involved in chloroplast proteolytic degradation, in orange proteins that form part of the chloroplast import machinery, in blue proteins involved in the UPS, and green circles represent Q28 or Q69 proteins.

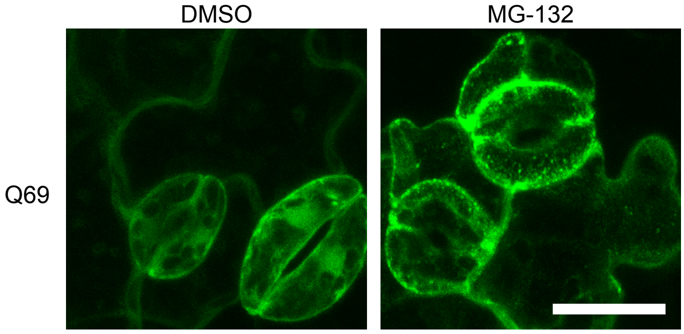


**Supplementary Figure 6. Q69 aggregates upon proteasome inhibition.** 7-day-old Q69 seedlings were transfer to liquid MS supplemented with DMSO or 50 μM MG-132. Images were taken after 8 hours of treatment. Scale indicates 20 μm.

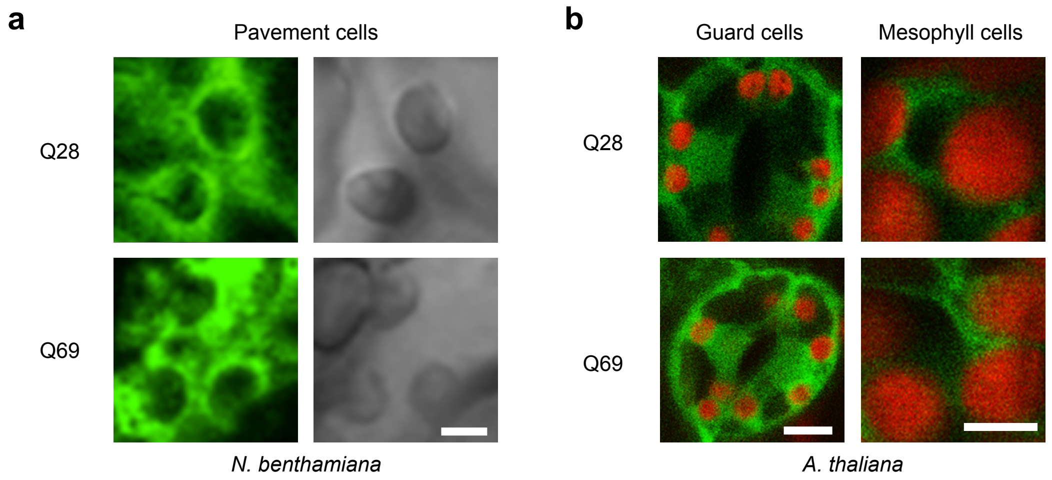


**Supplementary Figure 7. Q28 and Q69 accumulate around the chloroplasts. (a)** Q28 and Q69 distribution in *N. benthamiana* pavement cells from leaves analyzed at 6 days post-agroinfiltration (dpa). Citrine fluorescence and bright field showing the chloroplasts are shown. Scale indicates 5 µm. **(b)** Q28 and Q69 distribution in guard and mesophyll cells of 7-day-old transgenic Arabidopsis seedlings. Scales indicate 5 µm.

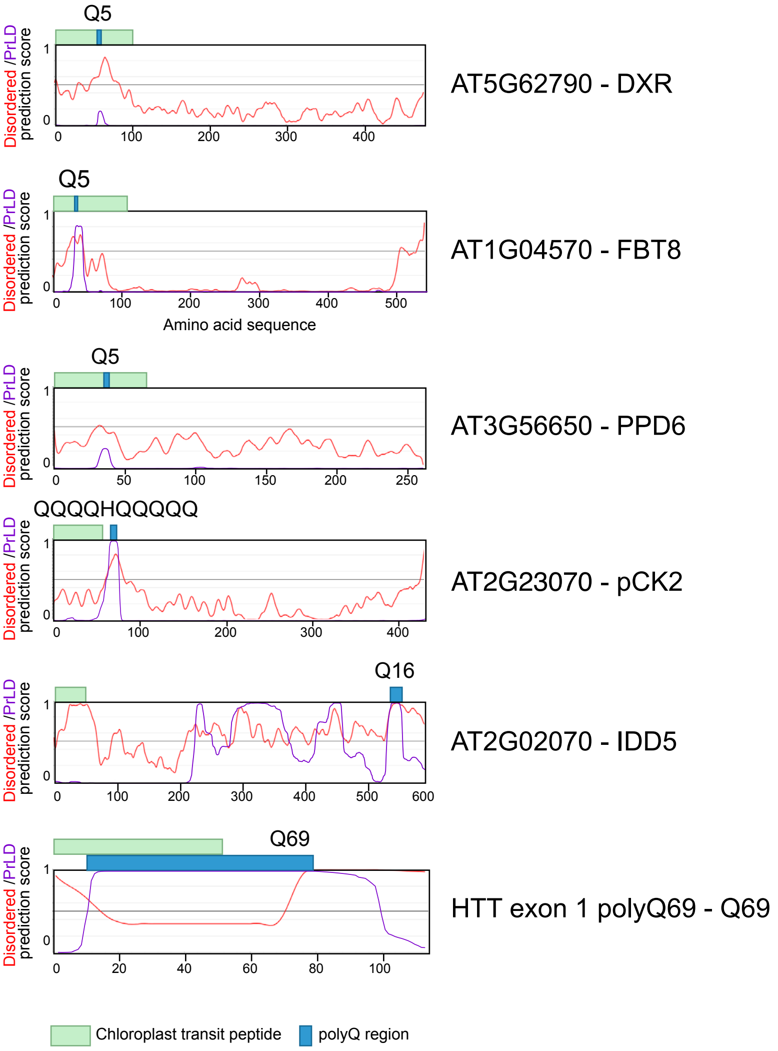


**Supplementary Figure 8. PolyQ regions are present in unstructured chloroplast transit peptides.** Prion-like amino score (purple line) was predicted using PLAAC (http://plaac.wi.mit.edu/). Protein unstructured score (red line) was predicted with IUPred3 (https://iupred.elte.hu). Green boxes represent the predicted transit peptide according to ChloroP 1.1. Blue boxes indicate the longest polyQ domain found in the amino acid sequence. Annotated chloroplast proteins were selected from Supplementary Data 1.

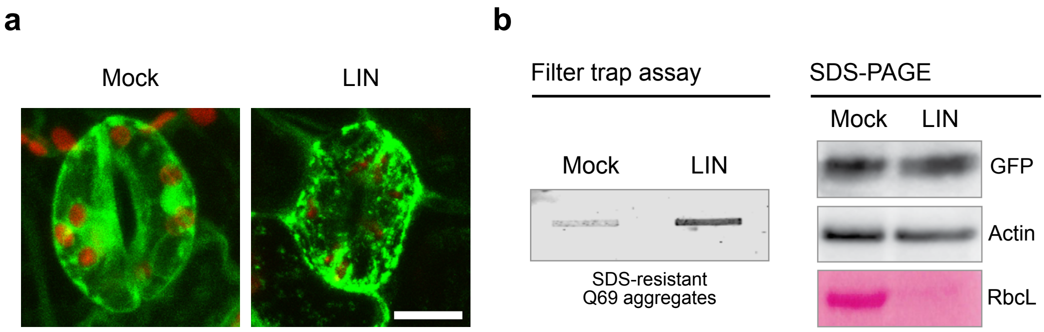


**Supplementary Figure 9. Q69 aggregates upon long-term LIN treatment. (a)** Representative confocal microscopy images showing Q69 distribution in 7-day-old seedlings grown and germinated in media supplemented with LIN 15 μM or mock solution. Images taken in stomata cells. Scale indicates 10 μm. **(b)** Filter trap and SDS-PAGE analysis of seedling from a. Representative blots of 3 independent experiments are shown.

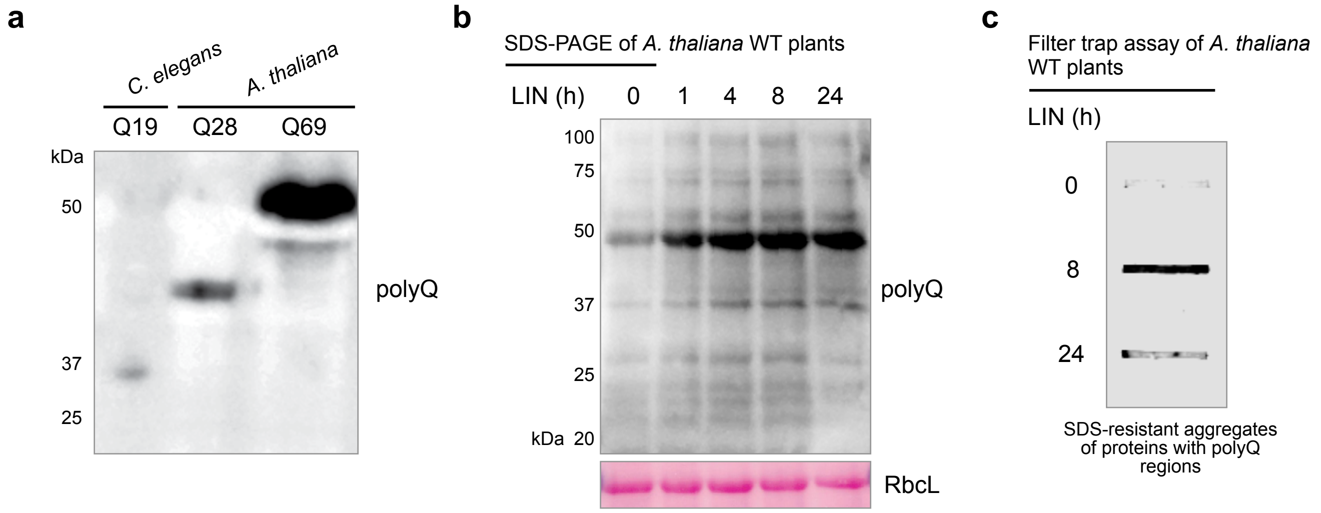


**Supplementary Figure 10. polyQ-containing Arabidopsis proteins accumulate upon Lin treatment.** **(a)** Immunoblot analysis of *Caenorhabditis elegans* expressing Q19, and constitutive transgenic Arabidopsis plants expressing Q28 and Q69. Immunoblot shows that polyQ antibody can recognize different size Q repeats. **(b)** Immunoblot analysis of 7-day-old WT plants transfer to liquid MS supplemented with 800 μM LIN. Samples were recollected at 0, 1, 4,8, and 24 hours (h) after LIN treatment. RbcL was used as loading control. Representative immunoblots of 3 independent experiments. **(c)** Filter trap analysis using polyQ antibody of protein samples of 7-day-old WT plants treated 0, 8 and 24 h with 800 μM LIN. Representative image of 3 experiments is shown.

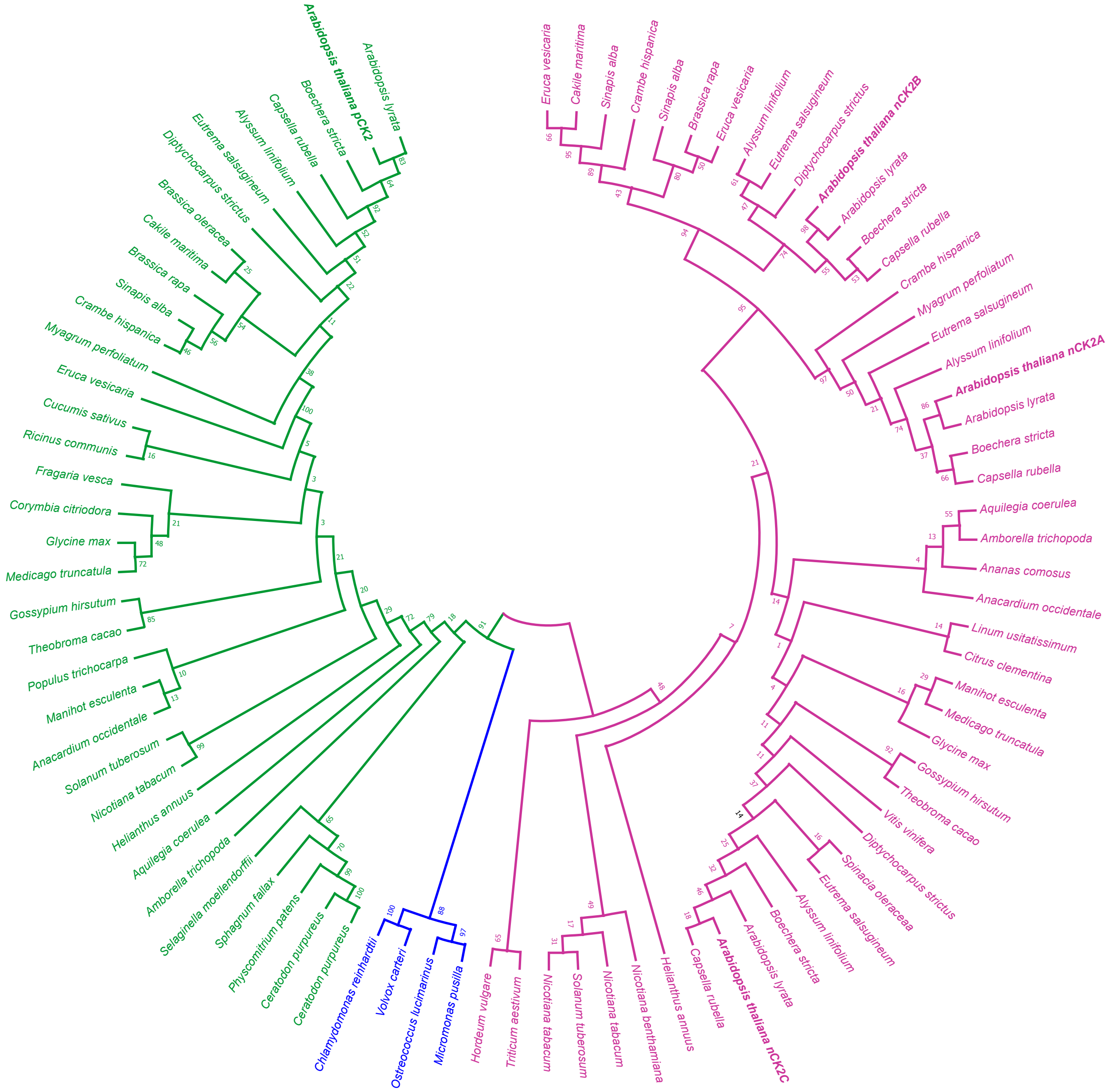


**Supplementary Figure 11. Plastid-specific CK2s cluster in a different clade than nuclear-specific CK2s in land plants.** Phylogenetic tree is based on the full-length sequences from the following photosynthetic organisms: *Arabidopsis lyrata* (AL4G12880), *Arabidopsis thaliana* (AT2G23070, pCK2), *Boechera stricta* (Bostr.21942s0016), *Capsella rubella* (Carub.0004s0227), *Alyssum linifolium* (Alyli.0381s0001), E*utrema salsugineum* (Thhalv10000167m), *Diptychocarpus strictus* (Distr.0027s16200), *Brassica oleracea* (Bol045814), *Cakile maritima* (Camar.1284s0005), *Brassica rapa* (Brara.I04552), *Sinapis alba* (Sialb.0006s0277), *Crambe hispanica* (Crahi.0145s0023), *Myagrum perfoliatum* (Myper.0012s0476), *Eruca vesicaria* (Eruve.0218s0002), *Cucumis sativus* (Cucsa.103080), *Ricinus communis* (29661.m000905), *Fragaria vesca* (FvH4_1g17410), *Corymbia citriodora* (Cocit.G1287), *Glycine max* (Glyma.17G161800), *Medicago truncatula* (Medtr4g095670), *Gossypium hirsutum* (Gohir.D11G207100), *Theobroma cacao* (Thecc.01G070500), *Populus trichocarpa* (Potri.007G054051), *Manihot esculenta* (Manes.13G112900), *Anacardium occidentale* (Anaoc.0017s0491), *Solanum tuberosum* (Soltu.DM.02G007610), *Nicotiana tabacum* (XP_016444753), *Helianthus annuus* (HanXRQChr14g0449481), *Aquilegia coerulea* (Aqcoe6G287100), *Amborella trichopoda* (evm_27.model.AmTr), *Selaginella moellendorffi*i (146942), *Sphagnum fallax* (Sphfalx16G059600), *Physcomitrium patens* (Pp3c2_21130V3), *Ceratodon purpureus* (CepurGG1.9G009900), *Ceratodon purpureus* (CepurR40.9G010400), *Chlamydomonas reinhardtii* (Cre14.g619166), *Volvox carteri* (Vocar.0004s0239), *Ostreococcus lucimarinus* (13142), *Micromonas pusilla* (148164), *Hordeum vulgare* (HORVU5Hr1G097230), *Triticum aestivum* (Traes_2DL_C1F0700CA), *Nicotiana tabacum* (CAD27342), *Solanum tuberosum* (Soltu.DM.03G005580), *Nicotiana_tabacum* (NP_001313201), *Nicotiana benthamiana* (AIT18344), *Helianthus annuus* (HanXRQChr14g0441851), *Capsella rubella* (Carub.0004s0228), *Arabidopsis thaliana* (AT2G23080, nCK2C), *Arabidopsis lyrata* (AL4G12890), *Boechera stricta* (Bostr.6864s0240), *Alyssum linifolium* (Alyli.0031s0202), *Eutrema salsugineum*

(Thhalv10010420m), *Spinacia oleracea* (Spov3_chr3.01021), *Diptychocarpus strictus* (Distr.0006s171500), *Vitis vinifera* (VIT_207s0129g00410), *Theobroma cacao* (Thecc.01G069000), *Gossypium hirsutum* (Gohir.A10G090200), *Glycine max* (Glyma.17G163700), *Medicago truncatula* (Medtr4g095400), *Manihot esculenta* (Manes.12G111700), *Citrus clementina* (Ciclev10025683m), *Linum usitatissimum* (Lus10019288), *Anacardium occidentale* (Anaoc.1135s0006), *Ananas comosus* (Aco012359), *Amborella trichopoda* (evm_27.model.AmTr), *Aquilegia coerulea* (Aqcoe5G073000), *Capsella rubella* (Carub.0005s1636), *Boechera stricta* (Bostr.0568s0162), *Arabidopsis lyrata* (AL8G43950), *Arabidopsis thaliana* (AT5G67380, nCK2A), *Alyssum linifolium* (Alyli.0048s0063), *Eutrema salsugineum* (Thhalv10000237m), *Myagrum perfoliatum* (Myper.0001s2551), *Crambe hispanica* (Crahi.0060s0084), *Capsella rubella* (Carub.0008s2711), *Boechera stricta* (Bostr.27895s0174), *Arabidopsis lyrata* (AL5G30060), *Arabidopsis thaliana* (AT3G50000, pCK2B), *Diptychocarpus strictus* (Distr.0027s16100), *Eutrema salsugineum* (Thhalv10004294m), *Alyssum linifolium* (Alyli.0171s0088), *Eruca vesicaria* (Eruve.0153s0024), *Brassica rapa* (Brara.I03327), *Sinapis alba* (Sialb.0274s0021), *Crambe hispanica* (Crahi.0183s0009), *Sinapis alba* (Sialb.0065s0203), *Cakile maritima* (Camar.4226s0003), and *Eruca vesicaria* (Eruve.2401s0010). The tree was constructed by rooting at midpoint using the maximum likelihood method with 2000 bootstrap replicates in MEGA11 (http://www.megasoftware.net) (Tamura et al. 2021). CK2 from Chlorophyte are displayed in blue. Plastid-specific CK2s (pCK2) and nuclear-specific CK2s (nCK2s) from Embryophyte are displayed in green and purple, respectively. Arabidopsis proteins are indicated in bold typeface.

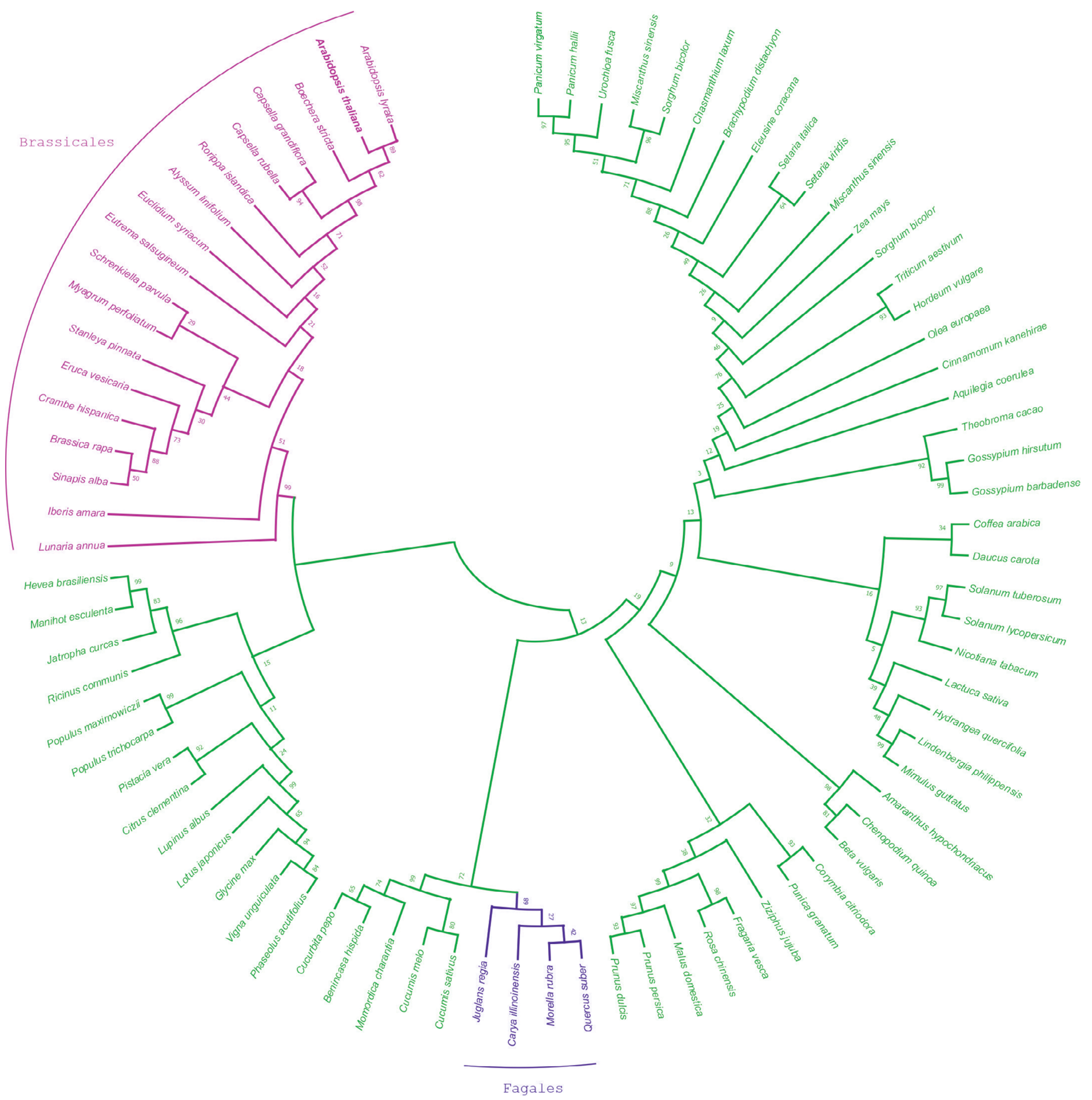


**Supplementary Figure 12. pCK2 polyQ domain is conserved in some dicotyledons such as Brassicales and Fagales.** Phylogenetic tree is based on the full-length sequences from the following plants: *Arabidopsis lyrata* (AL4G12880), *Arabidopsis thaliana* (AT2G23070), *Boechera stricta* (Bostr.21942s0016), *Capsella grandiflora* (Cagra.0068s0011), *Capsella rubella* (Carub.0008s2711), *Rorippa islandica* (Roisl.0046s0516), *Alyssum linifolium* (Alyli.0381s0001), *Euclidium syriacum* (Eusyr.0134s0504), *Eutrema salsugineum* (Thhalv10000167m), *Schrenkiella parvula* (Sp4g02090), *Myagrum perfoliatum* (Myper.0012s0476), *Stanleya pinnata* (Stapi.2640s0001), *Eruca vesicaria* (Eruve.0218s0002), *Crambe hispanica* (Crahi.0145s0023), *Brassica rapa* (Brara.I04552), ), *Sinapis alba* (Sialb.0006s0277), *Iberis amara* (Ibeam.1860s0015), *Lunaria annua* (Luann.0146s0054), *Hevea brasiliensis* (XP_021662590), *Manihot esculenta* (Manes.13G112900), *Jatropha curcas* (XP_012079387), *Ricinus communis* (29661.m000905), *Populus maximowiczii* (Poman.07G048700), *Populus trichocarpa* (Potri.007G054051), *Pistacia vera* (XP_031253414), *Citrus clementina* (Ciclev10025645m), *Lupinus albus* (Lalb_Chr16g0389201), *Lotus japonicus* (Lj4g0002812), *Glycine max* (Glyma.17G161800), *Vigna unguiculata* (Vigun03g285800), *Phaseolus acutifolius* (Phacu.CVR.003G282800), *Cucurbita pepo* (XP_023544339), *Benincasa hispida* (XP_038882656), *Momordica charantia* (XP_022132872), *Cucumis melo* (XP_008440369), *Cucumis sativus* (Cucsa.103080), *Juglans regia* (XP_035548062), *Carya illinoinensis* (Caril.02G004100), *Morella rubra* (KAB1214890), *Quercus suber* (XP_023870475), *Prunus dulcis* (XP_034219580), *Prunus persica* (Prupe.6G222700), *Malus domestica* (MD15G1295000), *Rosa chinensis* (XP_024181903), *Fragaria vesca* (FvH4_1g17410), *Ziziphus jujuba* (KAH7511196), *Punica granatum* (XP_031384638), *Corymbia citriodora* (Cocit.G1287), *Beta vulgaris* (EL10Ac6g15637), *Chenopodium quinoa* (XP_021714441), *Am-*

*aranthus hypochondriacus* (AH005694-RA), *Mimulus guttatus* (Migut.L00205), *Lindenbergia philippensis* (Liphi.04G118700), *Hydrangea quercifolia* (Hyque.13G027200), *Lactuca sativa* (Lsat_1_v5_gn_2_82920), *Nicotiana tabacum* (XP_016444753), *Solanum lycopersicum* (Solyc02g064700), *Solanum tuberosum* (Soltu.DM.02G007610), *Daucus carota* (DCAR_025788), *Coffea arabica* (evm.model.Scaffold_952.100), *Gossypium barbadense* (Gobar.D11G219600.1), *Gossypium hirsutum* (Gohir.D11G207100), *Theobroma cacao* (Thecc.01G070500), Aquilegia coerulea (Aqcoe6G287100), *Cinnamomum kanehirae* (CKAN_01168000), *Olea europaea* (Oeu048576), *Hordeum vulgare* (HORVU2Hr1G063420), *Triticum aestivum* (Traes_2DL_C1F0700CA), *Sorghum bicolor* (Sobic.002G010300), *Zea mays* (Zm00001d007951), *Miscanthus sinensis* (MisinT073100), *Setaria viridis* (Sevir.2G008300), *Setaria italica* (Seita.2G004600), *Eleusine coracana* (ELECO.r07.7AG0552810), *Brachypodium distachyon* (Bradi1g59010), *Chasmanthium laxum* (Chala.03G225300), *Sorghum bicolor* (Sobic.001G080700), *Miscanthus sinensis* (Misin01G066600), *Urochloa fusca* (Urofu.9G079300), *Panicum virgatum* (Pavir.9NG051900), and *Panicum hallii* (Pavir.9NG051900). The tree was constructed by rooting at midpoint using the maximum likelihood method with 2000 bootstrap replicates in MEGA11 (http://www.megasoftware.net) (Tamura et al. 2021). pCK2 from Brassicales and Fagales are displayed in purple and blue, respectively. Arabidopsis pCK2 protein is indicated in bold typeface.

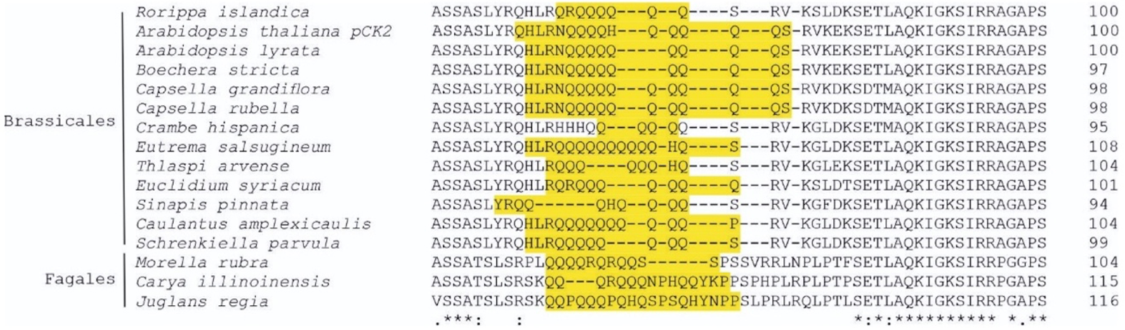


**Supplementary Figure 13. Prion-like domains are present in several pCK2 proteins in dicotyledons.** A partial alignment of the conserved polyQ at the N-terminal domain is shown. Alignment analysis was based on the full-length protein sequences from the following dicotyledon plants from Brassicales and Fagales orders: *Arabidopsis thaliana* (AT2G23070), *Arabidopsis lyrata* (AL4G12880), *Boechera stricta* (Bostr.21942s0016), *Capsella grandiflora* (Cagra.0068s0011), *Capsella rubella* (Carub.0004s0227), *Caulantus amplexicaulis* (Caamp.1039s0188), *Euclidium syriacum* (Eusyr.0134s0504), *Eutrema salsugineum* (Thhalv10000167m), *Rorippa islandica* (Roisl.0046s0516), *Schrenkiella parvula* (Sp4g02090), *Thlaspi arvense* (Thlar.0028s0002), *Sinapis pinnata* (Stapi.2640s0001), *Crambe hispanica* (Crahi.0145s0023), *Carya illinoinensis* (Caril.02G004100), *Juglans regia* (XP_035548062), and *Morella rubra* (KAB1214890). pCK2 sequences were identified using BLAST (https://blast.ncbi.nlm.nih.gov/Blast.cgi) and Phytozome (https://phytozome-next.jgi.doe.gov/), and they were aligned using Clustal Omega (https://www.ebi.ac.uk/Tools/msa/clustalo/). Prion-like domain residues were predicted using PLAAC (http://plaac.wi.mit.edu/) and are highlighted in yellow. Conserved residues are marked by asterisks.

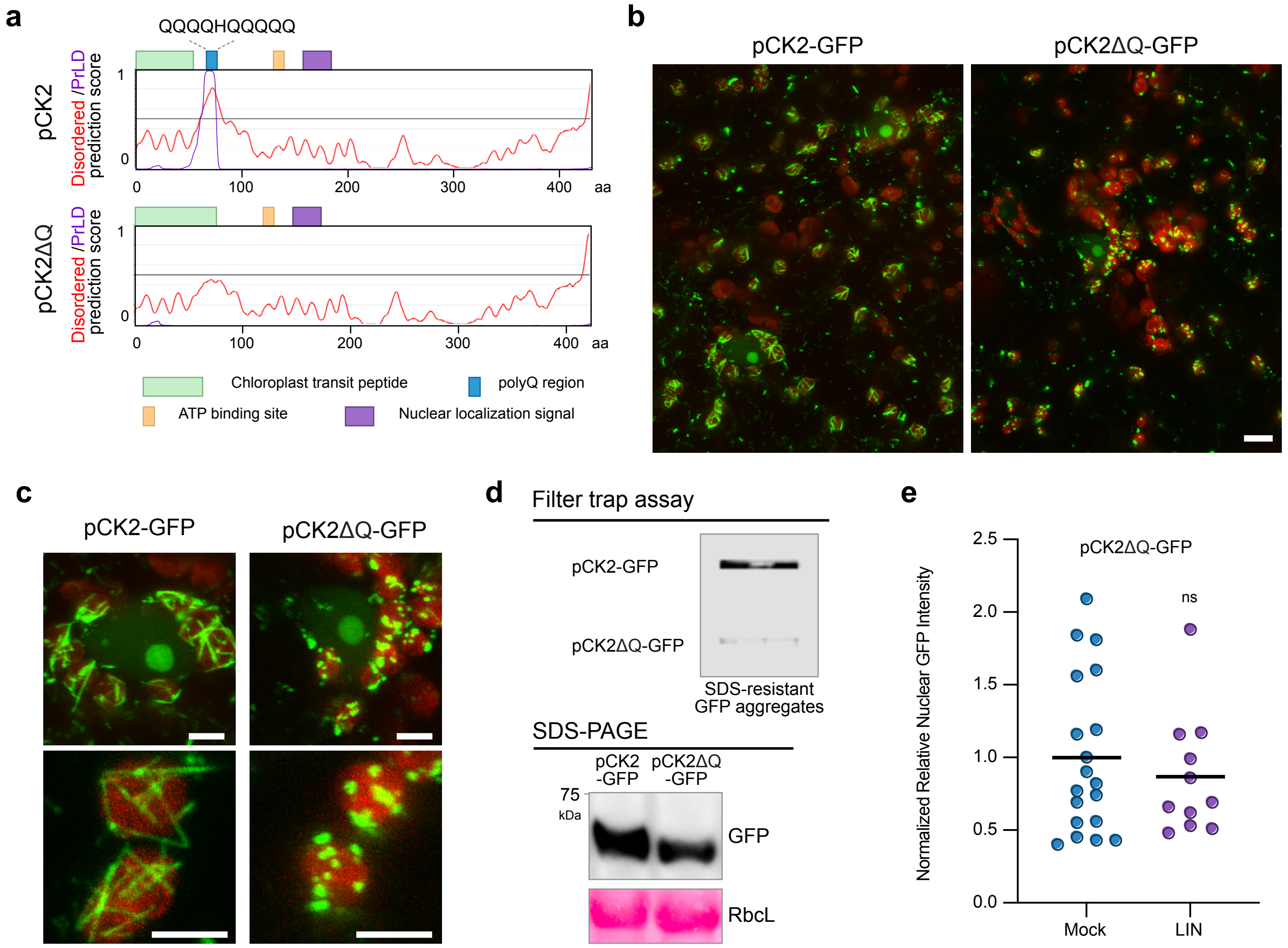


**Supplementary Figure 14. The polyQ region is necessary for the formation of pCK2 amyloid-like structures. (a)** Schematic representation of the construct used for pCK2 and pCK2ΔQ expression in *N. benthamiana* pavement cells. Scheme indicates position of different protein domains. Disordered (red line) and prion-like domain (purple line) prediction scores are also shown. **(b)** Confocal microscopy analysis of *N. benthamiana* pavement cells agroinfiltrated with 3*5S:pCK2-GFP* or *35S:pCK2ΔQ-GFP*. No amyloid-like structures were detected in plants agroinfiltrated with *35S:pCK2ΔQ-GFP*. Images are representative of 4 independent experiments. Scale indicates 10 μm **(c)** Closer magnification to nuclei and chloroplast of cells expressing pCK2-GFP or pCK2ΔQ. **(d)** Filter trap and SDS-PAGE analyses of *N. benthamiana* leaves expressing pCK2-GFP or pCK2ΔQ at 3 dpa. Representative immunoblots of 3 independent experiments are shown. **(e)** Graph representing quantification of nuclear GFP intensity of LIN or mock treated cells expressing pCK2ΔQ-GFP after 16 hours after infiltration of 800 μM LIN or mock solution. Leaves were infiltrated with LIN at day 3 after agroinfiltration. Mean and individual values for mock (n = 19) and LIN (n = 11) are shown.
